# Endothelial-immune crosstalk contributes to vasculopathy in non-alcoholic fatty liver disease

**DOI:** 10.1101/2021.12.13.472411

**Authors:** Chun Yi Ng, Khang Leng Lee, Mark D. Muthiah, Kan-Xing Wu, Florence W. J. Chioh, Konstanze Tan, Gwyneth S. T. Soon, Asim Shabbir, Wai Mun Loo, Zun Siong Low, Qingfeng Chen, Nguan Soon Tan, Huck Hui Ng, Yock Young Dan, Christine Cheung

## Abstract

The top cause of mortality in patients with non-alcoholic fatty liver disease (NAFLD) is cardiovascular complications. However, the mechanisms of NAFLD-associated vasculopathy remain understudied. We developed blood outgrowth endothelial cell (BOEC) models from NAFLD and healthy subjects. NAFLD BOECs exhibited global transcriptional upregulation of chemokine hallmarks and human leukocyte antigens. In mouse models of diet-induced NAFLD, we further confirmed enhanced endothelial expressions of CXCL12 in the aortas and liver vasculatures. To elucidate endothelial-immune crosstalk, we performed immunoprofiling by single-cell analysis, uncovering T cell intensification and potentially T-helper type 1 inflammation in NAFLD patients. Functionally, interference of the CXCL12-CXCR4 axis by small molecule AMD3100 selectively modulated the chemotaxis of patient-derived CD4+ T cells and natural killer cells towards NAFLD BOECs, restoring endothelial barrier integrity. Clinically, we detected three folds more circulating damaged endothelial cells in NAFLD patients than healthy controls. Our work provides insights for modulation of interactions with effector immune subsets to mitigate endothelial injury in NAFLD.

## Introduction

Non-alcoholic fatty liver disease is the most common liver disease in developed countries, affecting up to a third of western populations, and up to 40% in South East Asia (Muthiah and Sanyal, 2020). The spectrum of NAFLD includes the more benign non-alcoholic fatty liver, also known as simple steatosis, as well as the more progressive form non-alcoholic steatohepatitis (NASH) (Chalasani et al., 2018). NASH is characterized by lobular inflammation and ballooning, and can progress to develop fibrosis. While NAFLD remains a liver disease, the top cause of mortality in patients with NAFLD is cardiovascular mortality, which occurs independent of the shared metabolic risk factors such as insulin resistance and obesity (Targher et al., 2020). Patients with NAFLD are at increased risk of developing multiple vascular complications, including coronary artery disease, cerebrovascular disease, and peripheral vascular disease.

There is emerging evident on the link between NAFLD and impaired vascular health. Clinical measures of vascular function have established the association of reduced flow-mediated vasodilatation, altered carotid artery intimal medial thickness, coronary calcification and low coronary flow reserve with NAFLD (Federico et al., 2016, Sookoian and Pirola, 2008). In line with this, regression of NAFLD is associated with a lower risk of carotid atherosclerosis in observational cohort studies (Sinn et al., 2017, Sinn et al., 2016). Even in the absence of major cardiometabolic risk factors, healthy individuals demonstrated better vascular functions than NAFLD patients (Al-Hamoudi et al., 2020, Long et al., 2015), supporting that NAFLD alone could contribute to dysfunction of systemic vasculatures. Various mechanisms have been proposed as plausible explanations for accelerated atherosclerosis and increased cardiovascular risks in NAFLD patients, including a high oxidative stress state, macrophage activation (Targher et al., 2010) and systemic release of proatherogenic and thrombotic factors like tumor necrosis factor-α, interleukin-6, and oxidized low-density lipoprotein (Stols-Goncalves et al., 2019). Many of these paracrine mediators are known to activate endothelial cells and potentially result in endothelial dysfunction. Inflammation may also augment lipid risks to further drive atherosclerosis (Libby, 2021). However, there remain knowledge gaps in NAFLD-related endothelial disease biology that are not resolved by the current evaluation of vascular function using imaging modalities and serum biomarkers.

We aim to understand the molecular basis of endothelial pathophysiology in NAFLD-associated vasculopathy. Due to the difficulties of obtaining fresh vascular tissues from patients, we harnessed the use of blood outgrowth endothelial cells (BOECs) to develop personalized endothelial models from NAFLD patients and healthy subjects. BOECs originate from endothelial colony-forming cells which are bone marrow-derived progenitors found in the circulation and within vascular endothelium (Yoon et al., 2005). They are distinct from the early endothelial progenitor cells which form a heterogeneous culture of monocyte-derived cells from hematopoietic stem cell origin. Whereas BOECs give rise to a homogeneous population of mature endothelial cells with cobblestone morphology and are capable of tube formation. While the precursor of BOECs, endothelial colony-forming cells, have been explored for therapeutic revascularization, BOECs prove to be a useful surrogate of patient endothelial cells to investigate biology of vasculopathy in various disease contexts such as pulmonary arterial hypertension, diabetes and ischemic heart disease, etc (Paschalaki and Randi, 2018). Patient-derived BOECs were able to recapitulate clinical phenotypic alterations, capture the complexities of genetics and/or epigenetics, and demonstrate differential responses to disease-relevant stressors (Hebbel, 2017). Here, we utilized BOECs to facilitate experimentations including transcriptomic analysis, mechanistic interrogation and endothelial-immune cell coculture assays. We also analyzed circulating endothelial cells (CECs) which were shed from endothelial lining into the blood stream following vascular damage (Hebbel, 2017), to yield insights on endothelial injury in NAFLD.

## Results

### Development of human blood outgrowth endothelial cells for disease modelling

Patients with NAFLD were diagnosed with at least 5% steatosis characterized on liver biopsies, while selected healthy controls had a Controlled Attenuation Parameter score of less than 248 on vibration controlled transient elastography, indicative of less than 5% steatosis (more details in Materials and Methods under “patient enrolment”). We employed established protocols (Ormiston et al., 2015, Martin-Ramirez et al., 2012) to develop BOECs from NAFLD patients and healthy subjects. Briefly, peripheral blood mononuclear cells (PBMCs) isolated from whole blood samples were cultivated to derive BOECs (Fig. 1a). We then characterized the growth dynamics, marker expressions and functions of NAFLD and healthy control BOECs. BOEC colonies usually emerged after 7–14 days of PBMC cultivation (Fig. 1b). The colonies were isolated, passaged into endothelial growth media, and further cultured to stabilize the BOEC lines that displayed cobblestone endothelial cell morphology. Both NAFLD and healthy BOECs had comparable doubling times (Fig. 1c). More than 90% of BOECs expressed endothelial markers, including CD31 (PECAM1), CD144 (CDH5) and CD146, but negligible expressions for leukocyte markers CD45, CD14 and CD68, and progenitor cell marker CD133 (Fig. 1d), suggesting high purity of our BOEC derivation. Immunostaining confirmed their positive expressions of endothelial markers (Fig. 1e). Functionally, the BOECs were able to undergo angiogenesis, typical of endothelial property, in three-dimensional fibrin gel bead-based sprouting assays. Sprouts could be observed as early as 6h (Fig. 1f, representative images). An average of 8-10 sprouts per bead, with average sprout lengths of 100–150 µm were achieved by 24h (Fig. 1f, right panel). During sprouting angiogenesis, endothelial cells undergo a series of dynamic changes in their tip- and stalk-like cell phenotypes to facilitate cell migration, proliferation and stabilization, which ultimately leading to sprout formation (Blanco and Gerhardt, 2013). Indeed, our sprouted BOECs expressed significantly higher levels of tip- and stalk-like cell markers than monolayer BOECs (Supplementary Fig. S1). These quality control steps were performed to confirm mature endothelial phenotypes of both NAFLD and healthy control BOECs, before selected cell lines were used for downstream experimentations.

**Figure 1.**
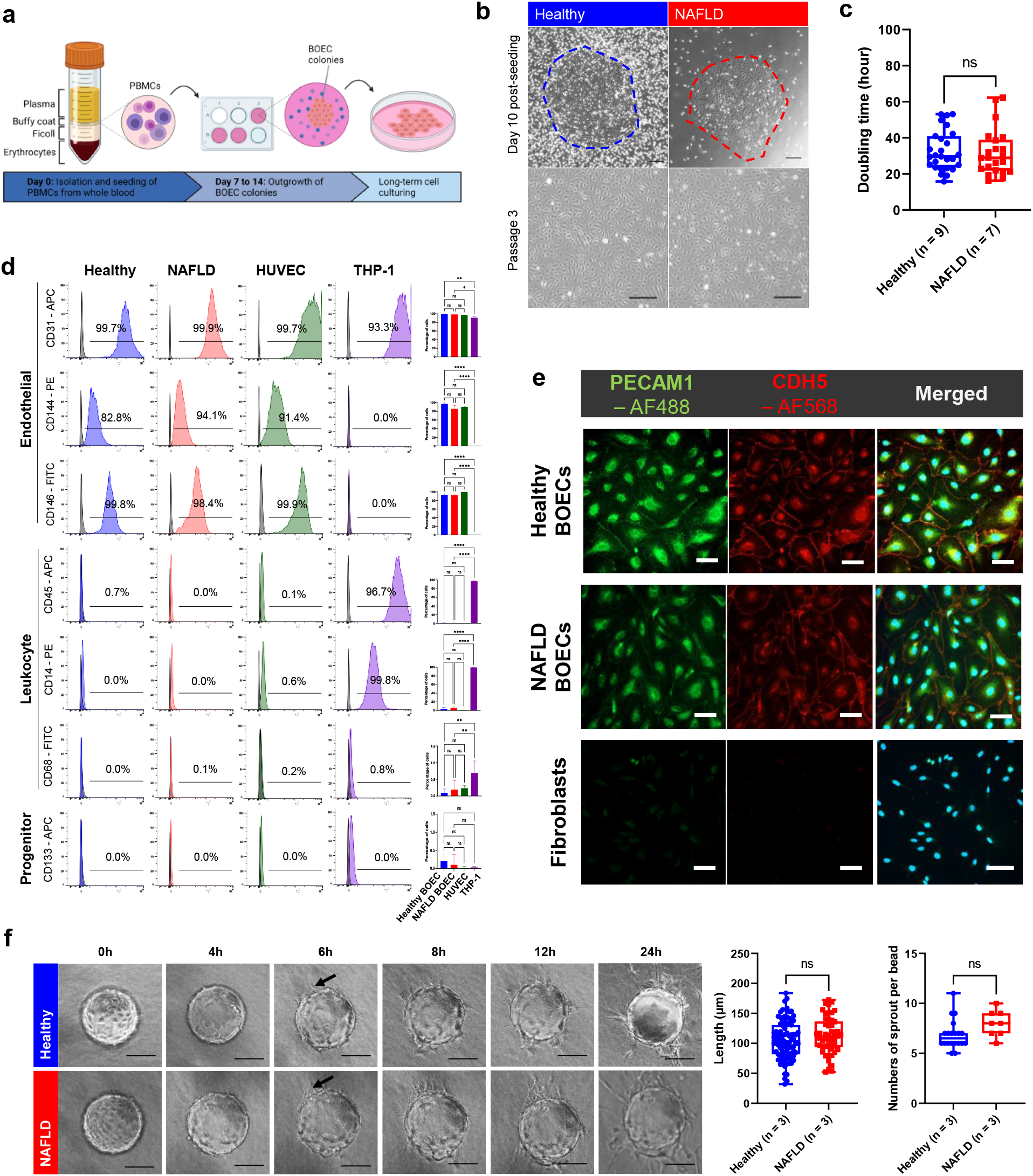
Generation and characterization of blood outgrowth endothelial cells from healthy and NAFLD subjects. **(a)** Workflow schematic of BOEC generation from PBMC samples isolated from NAFLD patients and healthy donors. **(b)** Top: Outgrowth of BOEC colonies from PBMC cultivation. Bottom: Stabilized BOEC cultures after passaging of colonies (scale bar, 250 µm). **(c)** Proliferation doubling time for NAFLD and healthy BOECs based on passages 2 and 3. Sample sizes are *n* = 9 healthy, *n* = 7 NAFLD. ns, not significant (*t*-test). **(d)** Flow cytometry characterization of BOECs for endothelial, leukocyte and progenitor cell markers (grey - isotype control; red/blue/green – cell lineage marker staining). HUVEC and monocytic THP-1 cells were used as positive controls for endothelial and leukocytic markers respectively. Right panel: Percentages of cells. Sample sizes are *n* = 4–6 healthy, *n* = 2–7 NAFLD, *n* = 1 fibroblasts. ns, not significant (one-way ANOVA). **(e)** Immunostaining of endothelial markers, PECAM1 and CDH5 (scale bar, 100 µm). **(f)** Longitudinal monitoring of BOEC angiogenesis in fibrin gel bead-based sprouting assays over 24h (representative images; scale bar, 100 µm). Right panel: Quantifications of sprout lengths and numbers of sprout per bead at 24h. *n* = 3. ns, not significant (*t*-test). All box plots indicate median (middle line), 25th, 75th percentile (box) and the lowest (respectively highest) data point (whiskers).

### NAFLD endothelial cells show enhanced chemokine hallmarks

To explore the molecular basis of endothelial pathophysiological mechanisms in NAFLD, we performed RNA-sequencing on BOECs derived from 3 NAFLD patients (2 females and 1 male, 37.7±11.5 years old) and 3 healthy controls (2 females and 1 male, 41.7±8.6 years old) (Supplemental Table S1). Principal component analysis (PCA) illustrated a clear segregation of transcriptomic profiles between NAFLD and healthy groups (Fig. 2a). A total of 670 differentially expressed genes were identified, including 535 upregulated genes and 135 downregulated genes in NAFLD BOECs in comparison to healthy BOECs (false discovery rate < 0.05, *p* < 0.05; fold change > 2 or < -2 respectively; Fig. 2b). Gene ontology analysis revealed that NAFLD BOECs are largely enriched in processes related to cell locomotion, extracellular matrix organization and chemotaxis, filament organization and mitogen-activated protein kinas (MAPK) cascade were most prominent in healthy BOECs (Fig. 2c). Based on the up-regulated genes in NAFLD BOECs, chemotaxis was identified as the most central network (Cluster 1) by molecular complex detection (MCODE), interconnecting with cell chemotaxis (leukocyte migration), cytoskeleton organization and cell morphogenesis (Fig. 2d). In contrast, no MCODE cluster was found using the down-regulated gene set (Supplemental Figure S2a), suggesting that the upregulated genes in NAFLD BOECs might exert a stronger biological effect due to intertwined processes. Correspondingly, heatmap visualization of the genes from the top enriched network showed that NAFLD BOECs expressed higher levels of chemokines including CC, CXC and CX3C families than healthy BOECs (Fig. 2e).

**Figure 2.**
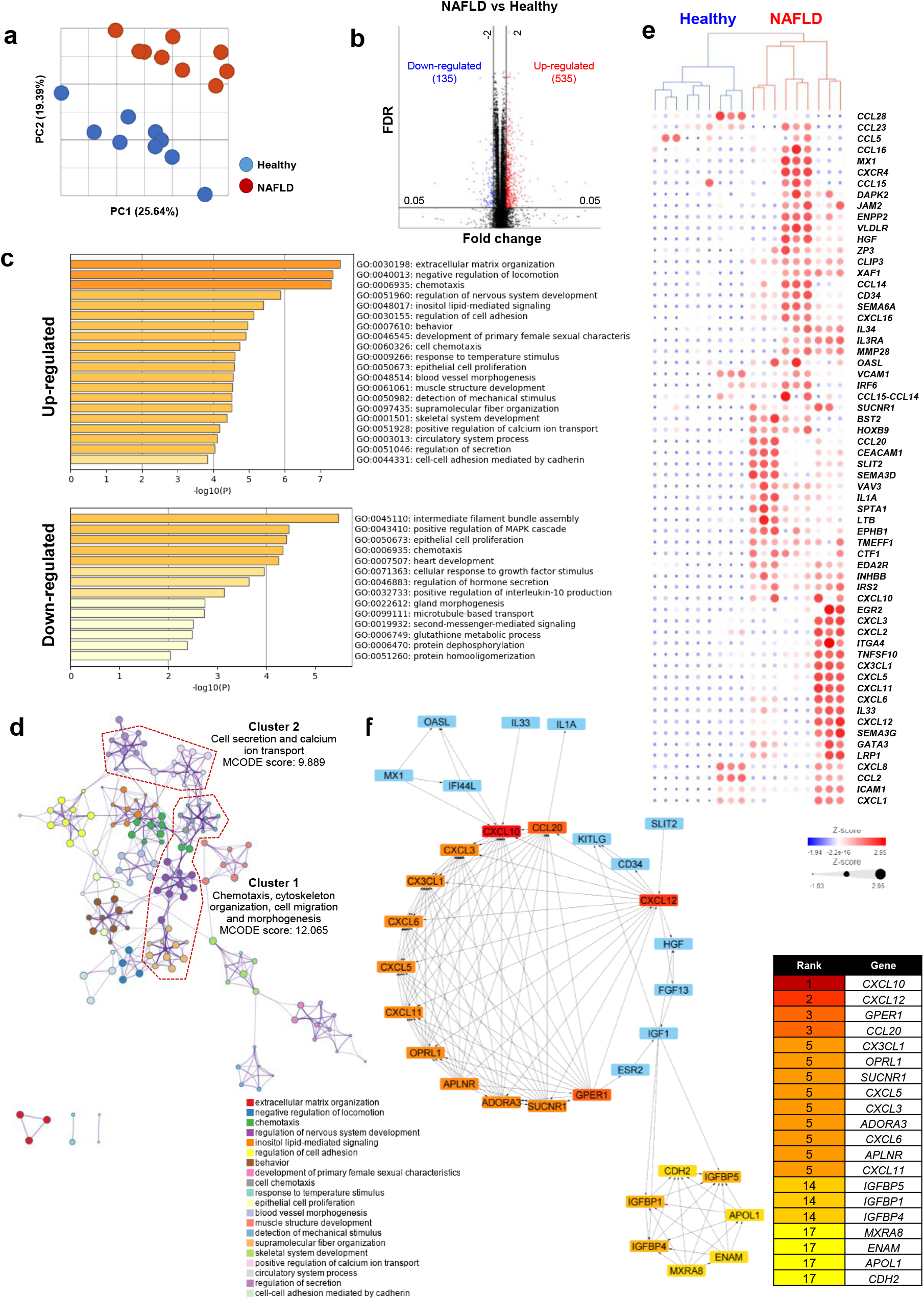
Transcriptomic analysis of NAFLD and healthy BOECs. **(a)** Principal component analysis (PCA) of NAFLD (*n* = 3) and healthy (*n* = 3) BOEC transcriptomes. **(b)** Volcano plot depicting differentially expressed genes comparing NAFLD BOECs with healthy BOECs. Red and blue dots represent upregulated and downregulated genes in NAFLD BOECs, respectively. **(c)** Enriched GO terms of the differentially expressed genes. **(d)** Metascape enrichment network visualization showing intra- and inter-cluster interactomes of the upregulated genes in NAFLD BOECs. Nodes that share the same cluster annotations are close to each other. Densely connected clusters were identified using MCODE algorithm and outlined in red. **(e)** Heatmap featuring the genes in MCODE Cluster 1. **(f)** STRING protein-protein interaction network were analyzed by Maximum Neighborhood Component, showing hub genes as ranked and indicated by gradient of red colors. Blue nodes represent differentially expressed genes in the extended networks.

To prioritize candidate genes for further mechanistic interrogation, we used 12 topological data analysis methods to identify and rank top 20 genes per method which might play a central role in the network interactome (Supplemental Table S2). Chemokines including *CXCL3, CXCL5, CXCL6, CXCL10, CXCL11, CXCL12, CCL20, and CX3CL1* were consistently reflected by at least 6 topological analysis methods. Of those, *CXCL10, CXCL12 and CX3CL1* were ranked as top 3 genes by at least three methods, emphasizing their importance in the network. Fig. 2f is a visualization of one of the methods used. We also applied MCODE algorithm that verified ‘chemokine-mediated signaling pathway’ as the top MCODE complex containing most of the aforementioned chemokine genes (Supplemental Fig. S2b). Taken together, enhanced chemokine signatures could be a hallmark for NAFLD endothelial cells.

### NAFLD patient endothelial cells and vascular endothelia of in vivo disease models demonstrate intensified CXCL12 levels

We selected several chemokines from the prioritized list of candidates for further validation in a larger number of endothelial cell lines. Gene expression profiling demonstrated that *CXCL10* and *CXCL12* were significantly upregulated in NAFLD BOECs (*n* = 11) compared to healthy BOECs (*n* = 12), while the other chemokines were not significantly different at baseline (Fig. 3a, donor demographics in Supplemental Table S3). To examine these chemokine expressions under relevant pathophysiological context, we introduced autologous plasma as a stressor to the BOEC cultures. Upon exposure of BOECs to their autologous plasma, similarly, CXC chemokine ligands were significantly upregulated in NAFLD BOECs than in healthy BOECs (Supplemental Fig. S3a). It was reported that levels of free fatty acids (FFAs) and lipopolysaccharide (LPS) in plasma were positively correlated with NAFLD (Zhang et al., 2014, Zhu et al., 2015). These factors are known to activate endothelial cells and exacerbate inflammation. We further examined the chemokine expression profile of BOECs in the presence of FFAs and LPS. There were no significant differences in lipid uptake by both NAFLD and healthy BOECs (Fig. 3b). Under exposure to FFAs and LPS, NAFLD BOECs had visibly more pronounced *CXCL12* and *CX3CL1* expressions than healthy controls (Fig. 3c). NAFLD BOECs seemed intrinsically primed with higher *CXCL12* expression than healthy BOECs.

**Figure 3.**
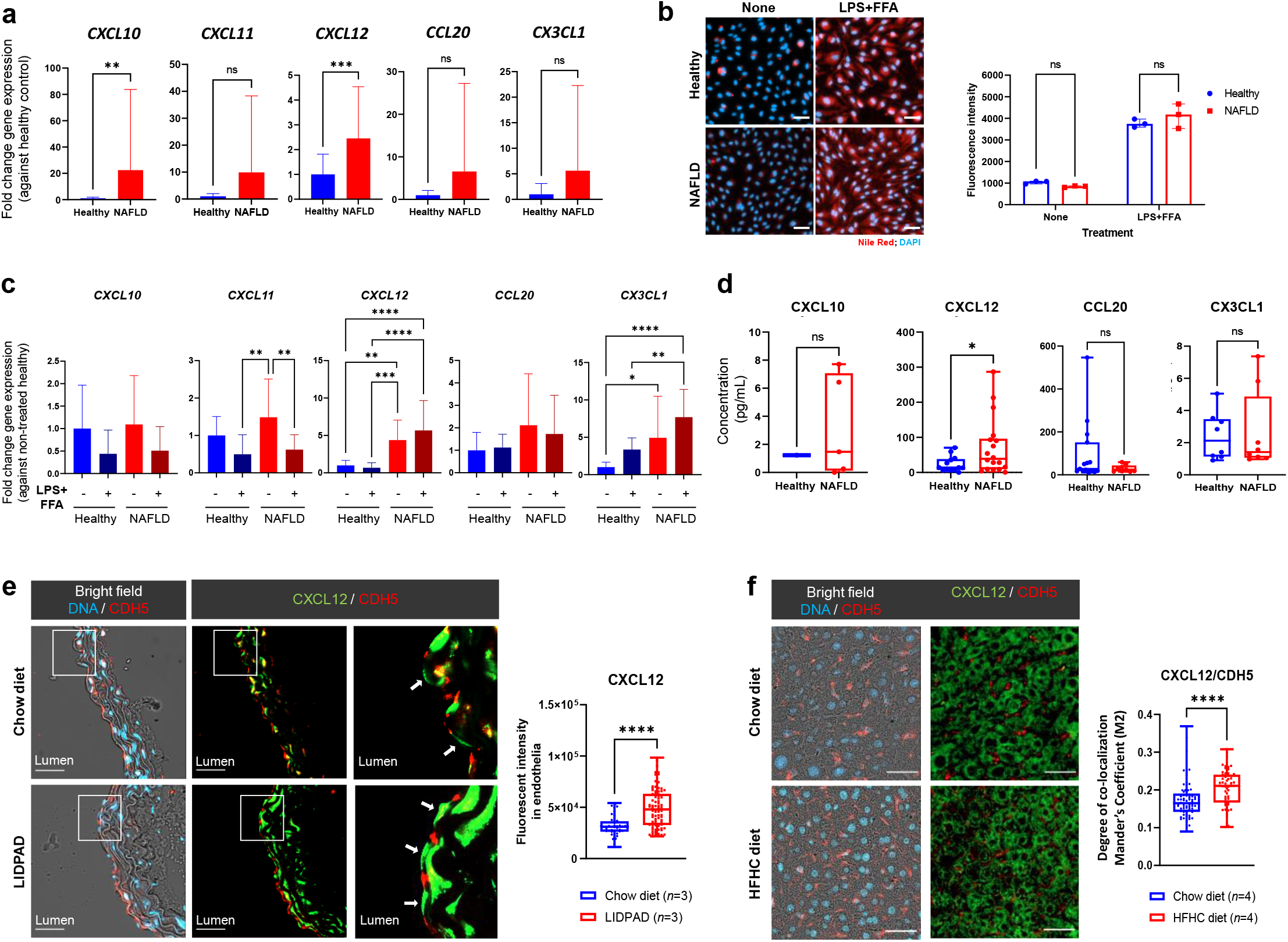
Validation of chemokine hallmarks in human endothelial cells and mouse disease models. **(a)** Fold change in gene expressions of *CXCL10, CXCL11, CXCL12, CCL20* and *CX3CL1* in NAFLD (*n* = 11) BOECs, normalized to healthy (*n* = 12) BOECs. *ns*, not significant; \*\*\**p*<0.001 (Mann-Whitney test). Results are indicated by mean ± SD. **(b)** Left: Representative images showing non-treated and LPS+FFA-treated NAFLD and healthy BOECs, stained with Nile Red (scale bars, 100 µm). Right: Fluorescence intensity of intracellular Nile Red stain was quantified at 515/585nm, and readings were normalized with blanks (*n =* 3). *ns*, not significant, *p>*0.05 (Two-way ANOVA). Results are indicated by median with 95% CI. **(c)** Fold change in gene expressions of *CXCL10, CXCL11, CXCL12, CCL20* and *CX3CL1* in LPS+FFA- or non-treated NAFLD (*n* = 5) BOECs, normalized to non-treated healthy (*n* = 4) BOECs. ns, not significant; \*\*\*\**p*<0.0001 (Two-way ANOVA). Results are indicated by mean ± SD. **(d)** Concentrations of chemokines in conditioned culture media of healthy and NAFLD BOECs. ns, not significant; \**p*<0.05 (*t*-test). Box plots indicate median (middle line), 25th, 75th percentile (box) and the lowest (respectively highest) data point (whiskers). **(e)** Top: Representative images (BF, bright field; DAPI, nuclei; CDH5, endothelial cells; CXCL12) of aortic sections from chow diet-fed (*n* = 3) and LIDPAD (*n* = 3) mice (scale bar, 40 μm). Bottom: Box-whisker plot of CXCL12 fluorescent intensity in the aortic vascular endothelia. CXCL12 fluorescent intensity for each delimitated endothelial region is represented as individual data points on the plot, with horizontal bars indicating the mean, minimum and maximum values for each group. \*\*\*\**p*<0.0001 (Mann-Whitney test). **(f)** Top: Representative images (BF, bright field; DAPI, nuclei; CDH5, endothelial cells; CXCL12) of liver sections from chow diet-fed (*n* = 4) and HFHC diet-fed (*n* = 4) humanized mice (scale bar, 40 μm). Bottom: Box-whisker plot of Mander’s Coefficient (M2) that was a measure of the degree of CXCL12 overlap with CDH5-positive vasculatures in the livers. \*\*\*\**p*<0.0001 (Mann-Whitney test). There were 10-20 independent regions of interest analyzed per mouse for image quantification in (c) and (d).

Previous studies have also shown that serum CXCL10 and CXCL12 levels were statistically higher by severity of fibrosis, steatosis and inflammation (Chalin et al., 2018). Our ELISA quantification of these chemokines in plasma samples, although insignificant, showed a consistent trend with higher CXCL10 and CXCL12 levels in NAFLD plasma than healthy controls (Supplemental Fig. S3b). Then we tested if NAFLD endothelial cells could be a contributor of soluble chemokines. Our ELISA data confirmed an intensified secretion of CXCL12 protein in NAFLD BOECs-conditioned culture media, after correction against non-conditioned media (Fig. 3d).

We decided to bring forward CXCL12 as the key candidate to validate in two murine NAFLD models. An improved diet-inducible liver disease mouse model was established by the use of a new modified LIDPAD (Liver Disease Progression Aggravation Diet) diet (Low et al. manuscript in submission elsewhere). As systemic vasculopathies may be associated with NAFLD, we analyzed aortic tissue sections harvested from mice after 12 weeks of LIDPAD or control diets. Histological staining showed that CXCL12 was ubiquitously expressed throughout the aortic intima and media (representative images, Fig. 3e). We identified CDH5-positive endothelial cells which showed intercellular junctional staining pattern (CDH5 in red, Fig. 3e). Quantitatively, CXCL12 expressions within regions of interest strictly containing just CDH5-positive endothelial lining were found to be significantly higher in LIDPAD aortic sections compared with chow diet group. In addition, we examined liver sections harvested from humanized mice that were reconstituted with human immune components (Her et al., 2020b). The high fat and high calorie (HFHC) diet induces development of liver steatosis, accompanied by inflammation and fibrosis, which are characteristic features of human advanced NAFLD (Her et al., 2020b). Humanized mice on a 20-week HFHC diet demonstrated higher levels of CXCL12 associated with the liver vasculatures as compared to mice on chow diet (Fig. 3f). Notably, in our previous study, T lymphocyte infiltration into liver was apparent in mice on HFHC diet (Her et al., 2020b). Hence, we were motivated to find out if NAFLD endothelial chemokines could play a role in regulating crosstalk with immune cells.

### NAFLD endothelial cells may preferentially interact with effector lymphocytes

Interplay of endothelial cells and leukocytes implicates vascular inflammation and atherogenesis (Libby, 2021). We hypothesized that analysis of chemokine receptor expressions in patient immune populations would allow us to better predict immune interactors of vascular endothelia in NAFLD. Using single-cell RNA-sequencing, we performed immunoprofiling on peripheral blood mononuclear cell (PBMCs) samples from 2 NAFLD patients (1 make, 1 female) and 4 healthy controls (2 males, 2 females) (Supplemental Table S4). In the UMAP visualization, we identified 14 distinct cell type clusters by DICE annotation (Schmiedel et al., 2018) (top panel, Fig. 4a). Based on chemokine ligand-receptor mapping (Hughes and Nibbs, 2018), we analyzed expressions of the counter receptors (lower panel, Fig. 4a) to those chemokine candidates enriched in NAFLD endothelial cells. Most of the chemokine receptor genes showed differential expressions across various immune populations, while *CXCR4* was more ubiquitously expressed. Chemokine receptor-expressing populations were primarily found in T lymphocytes, natural killer (NK) cells and monocytes (Fig. 4b). Within the T lymphocytes, chemokine receptors were more prominently expressed by CD4+ effector T helper (Th) subsets, Th1 and Th17 cells, which are known to secret proinflammatory cytokines such as interferon-γ and interleukin-17 (Leung et al., 2010). Correspondingly, we observed an overarching T cell intensification in NAFLD but diminished proportions of NK cells and monocytes (Fig. 4c). In both NAFLD and healthy groups, Th1 cells formed the largest subset among effector T helper populations, motivating us to perform gene ontology enrichment analysis on their differentially expressed genes in Th1 cell subset. Various aspects of transcription and translation regulations were common to both NAFLD and healthy groups (Fig. 4d). Th1 cells from healthy controls were enriched in regulation of immune response. Notably, NAFLD Th1 subset seemed more proinflammatory in nature by having enriched processes relating to production of interferons and tumor necrosis factor, as well as NF-κB signaling (Fig. 4d).

**Figure 4.**
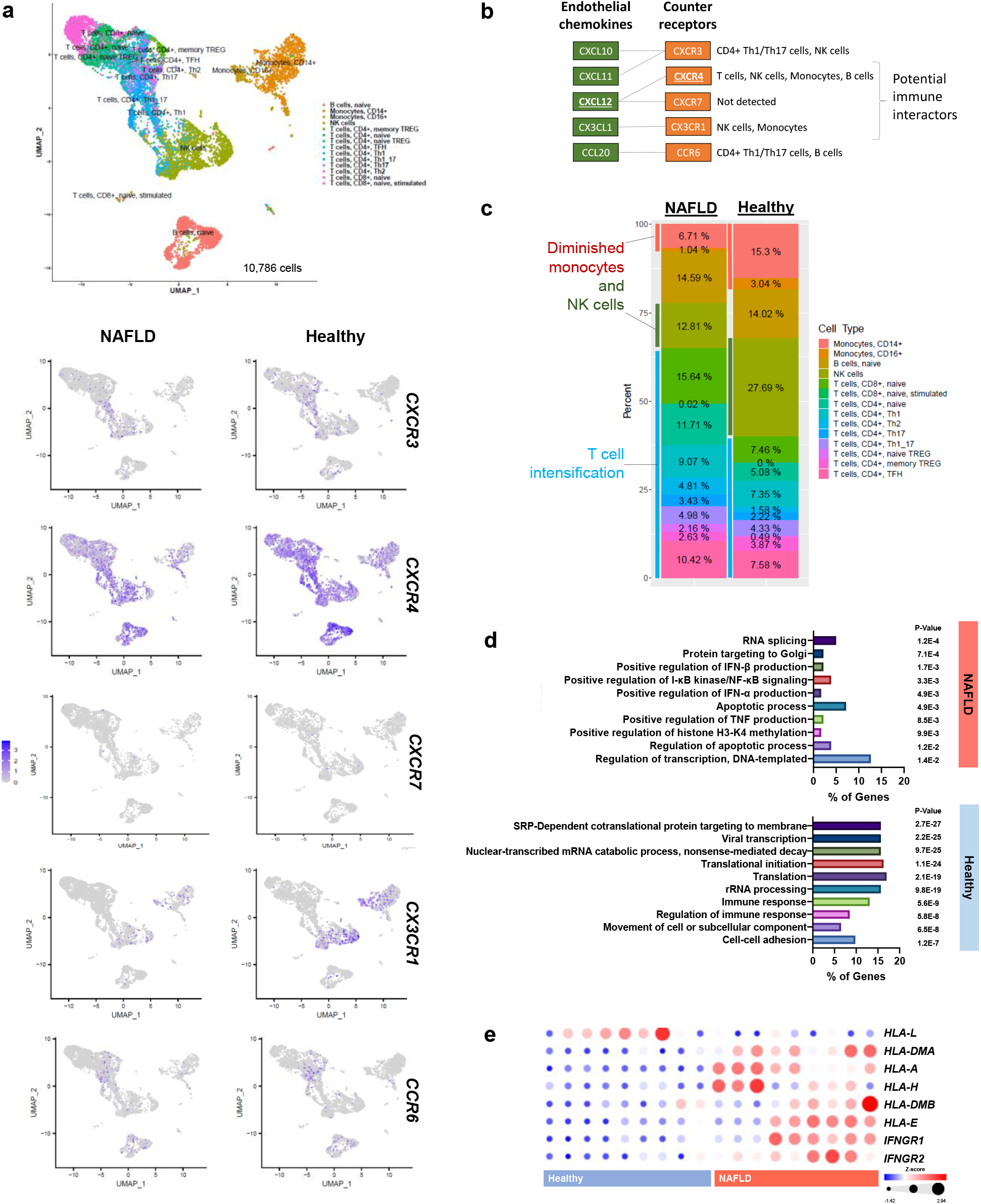
Immunoprofiling of NAFLD patients and healthy subjects by single-cell transcriptomics. **(a)** Top: Single-cell UMAP of sequenced PBMCs from 2 NAFLD patients and 4 healthy subjects; *n* = 10,786 cells. We identified 15 distinct cell types by DICE annotation. Bottom: Expression patterns of *CXCR3, CXCR4, CXCR7, CX3CR1, CCR6* across different cell types in NAFLD and healthy samples. **(b)** Chemokine ligand-receptor mapping to identify potential immune interactors based on NAFLD endothelial chemokine signatures. **(c)** Graphic shows the percentage composition of identified cell types in NAFLD and healthy PBMCs single-cell analysis. **(d)** Top 10 gene ontology biological processes of T helper 1 subset in NAFLD (188 differentially regulated genes) and healthy (157 differentially regulated genes), based on logFC≥0.3, p<0.001. Analysis was done using the Database for Annotation, Visualization and Integrated Discovery (DAVID) v6.8. **(e)** Heatmap of relevant T lymphocyte-interacting genes expressed in NAFLD and healthy BOECs.

To decipher if NAFLD endothelial cells could potentially interact with T lymphocytes, we retrieved relevant gene expressions from the BOEC transcriptomic dataset, particularly the human leukocyte antigen (HLA) family members. HLA Class 1 members mainly support interaction with CD8+ T cells, whereby HLA Class 2 members facilitate binding with CD4+ T subsets (Laidlaw et al., 2016). We found significantly higher expressions of several HLAs in NAFLD BOECs than healthy BOECs, involving Class 1 members *HLA-A, HLA-E* and *HLA-H*, and Class 2 members *HLA-DMA* and *HLA-DMB* (Fig. 4e). Interestingly, NAFLD BOECs also had significantly higher expressions of interferon-γ receptors *IFNGR1* and *IFNGR2* (Fig. 4e), potentially making them susceptible to interferon-γ that can modify activated phenotype of endothelial cells to favor Th1 cell inflammatory reactions (Pober and Sessa, 2007). The data thus far suggest that endothelial cells in NAFLD may be providing a chemokine milieu and activated phenotype that preferentially support the interaction with effector T cells.

### CXCL12-CXCR4 axis is implicated in NAFLD endothelial-immune crosstalk

Phenotypically, to study the interaction of NAFLD endothelial cells with immune mediators, we performed coculture assays of endothelial-immune cell adhesion, chemotaxis and transendothelial migration. We first employed two types of human leukocyte cell lines – T lymphocytic Jurkat cells and monocytic THP-1 cells for adhesion assay. Both NAFLD and healthy BOECs demonstrated greater extent of adhesion with Jurkat cells than THP-1 cells onto their monolayers (Fig. 5a). In particular, significantly more Jurkat cells adhered to NAFLD BOECs than to healthy BOECs. Since NAFLD BOECs were found to be activated with chemokine hallmarks (Fig. 3), we further confirmed that NAFLD BOECs had significantly greater chemotactic ability than healthy BOECs in recruiting Jurkat cells, but THP-1 cells (Fig. 5b). These data seemed to indicate preferential recruitment of T lymphocytes to NAFLD BOECs.

**Figure 5.**
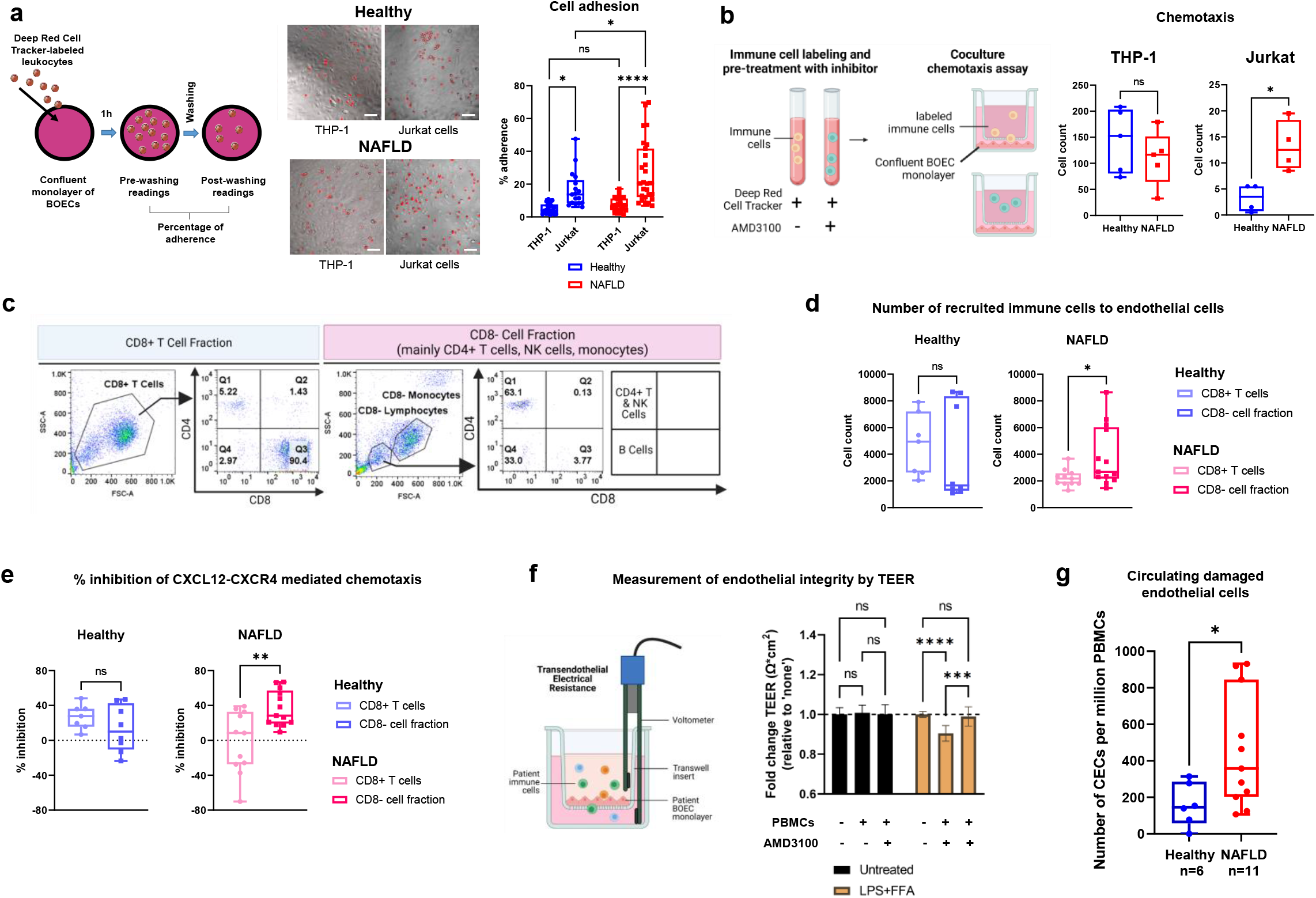
Characterizations of endothelial-immune interactions. **(a)** Left: Workflow schematic of adhesion assay. Middle: Representative images of labeled THP-1 and Jurkat cells attached to BOEC monolayers (Scale bars, 100 µm). Right: Scatter box plots of the numbers of adhered leukocytes were determined fluorometrically. Sample size is *n* = 9 NAFLD, *n* = 7 healthy; \**p* < 0.05; \*\*\*\**p* < 0.0001; *ns*, non-significant (two-way ANOVA). **(b)** Left: Schematic of chemotaxis assay. Right: Scatter box plots of the numbers of recruited THP-1 and Jurkat cells by the BOECs. Sample size is *n* = 5; \**p* < 0.05; *ns*, non-significant (*t*-test). **(c)** Representative flow cytometry dot plots, characterizing the isolated CD8+ T cells and CD8-immune cell fraction (mainly CD4+ T cells, NK cells and monocytes) from PBMCs of healthy and NAFLD subjects. **(d)** Numbers of recruited immune cells to BOECs in coculture chemotaxis assay. Sample size is *n* = 4 healthy; *n* = 6 NAFLD. \**p* < 0.05; *ns*, non-significant (*t*-test). **(e)** Percentages of inhibition on chemotaxis after pre-treatment with AMD3100. Sample size is *n* = 3–4 healthy; *n* = 5–6 NAFLD. \*\**p* < 0.01; *ns*, non-significant (*t*-test). **(f)** Left: Schematic of transendothelial migration assay with patient-derived BOECs and immune cells. Right: Transendothelial electric resistance (TEER) measurements of BOEC monolayers in the presence and absence of PBMCs and/or AMD3100. Sample size is *n* = 3. \*\*\**p* < 0.001; \*\*\*\**p* < 0.0001; *ns*, non-significant (two-way ANOVA). **(g)** Scatter box plots of the numbers of circulating endothelial cells per million PBMCs in healthy (*n* = 6) and NAFLD subjects (*n* = 11). \**p* < 0.05 (Mann-Whitney test). All box plots in this figure indicate median (middle line), 25th, 75th percentile (box) and the lowest (respectively highest) data points (whiskers).

To better mimic pathophysiological context, we switched to using PBMCs obtained from NAFLD patients and healthy subjects for further coculture studies with BOECs. As the chemokine receptor-expressing populations mainly involved CD4+ T cells, NK cells and monocytes (Fig. 4), we conducted magnetic-activated immune cell sorting to derive CD8+ and CD8-(largely include CD4+ effector T cells, NK cells, monocytes) cell fractions from NAFLD and healthy PBMCs, followed by flow cytometry characterization (Fig. 5c). Functionally, we assessed the chemotactic migration of NAFLD and healthy CD8+/-immune cell fractions toward BOEC monolayers matched by their health conditions. Interestingly, we observed preferential chemotaxis of NAFLD CD8-cell fraction (primarily CD4+ T cells, NK cells and monocytes) in coculture with NAFLD BOECs, but not in the healthy control coculture setup (Fig. 5d). The decreased ability of NAFLD BOECs in recruiting CD8+ immune cells was in consensus with our immunoprofiling findings that the chemokine receptor-expressing populations primarily converge on CD4+ T cells, NK cells and monocytes. Since CXCL12 was consistently enhanced in NAFLD patient endothelial cells, as well as in vascular endothelia of *in vivo* disease models (Fig. 3), we postulated the involvement of CXCL12-CXCR4 axis in the preferential recruitment of CD4+ effector T cells and NK cells in NAFLD. To intervene the CXCL12-CXCR4 axis in endothelial-immune chemotaxis, AMD3100, a small molecule inhibitor of CXCR4, was chosen as it has been validated and approved for human use (Hatse et al., 2002, Fricker et al., 2006). Specificity of AMD3100 is attributed to its well-defined high-affinity molecular interactions between the cyclam moieties of drug molecule and aspartate residues of CXCR4 receptor (Gerlach et al., 2001). Here, we found that chemotactic migration of CD8-cell fraction was attenuated significantly in the presence of AMD3100 in NAFLD coculture assay but not in the healthy group (Fig. 5e). These findings suggested differences between NAFLD and healthy control in the recruitment of distinct immune subsets to endothelial cells, which could be mediated through the CXCL12-CXCR4 axis.

To understand the vascular consequences of endothelial-immune crosstalk, we characterized NAFLD BOEC barrier integrity after transendothelial migration assay with NAFLD PBMCs. We measured transendothelial electrical resistance (TEER) of BOEC monolayers, where lower TEER readings would indicate greater endothelial barrier permeability. At baseline (Fig. 5f, Untreated group), we did not observe changes of TEER upon transmigration of NAFLD PBMCs. However, perturbations of endothelial barrier became apparent when NAFLD BOECs were pre-treated with LPS and FFA, as seen in the significant reduction of TEER readings after interaction with NAFLD PBMCs (Fig. 5f, LPS+FFA group). Both LPS and FFA are factors known to be implicated in NAFLD, rendering BOECs more sensitive to immune-inflicted injury. Inhibition of CXCL12-CXCR4 axis by AMD3100 averted the impact of PBMC transmigration and restored TEER readings (Fig. 5f, LPS+FFA group). Taking these findings to clinical setting, we analyzed circulating endothelial cells (CECs) that were cells shed from damaged blood vessels into the bloodstream, hence indicative of the level of vascular injury (Hebbel, 2017). NAFLD patients had more than 3 folds of CECs compared to healthy subjects (Fig. 5g), validating that endothelial pathology was indeed aggravated under NAFLD condition.

## Discussion

With the prevalence of NAFLD, there is an increasing recognition of cardiovascular complications being significant causes of mortality in these patients. In this study, we have advanced mechanistic understanding of endothelial pathophysiology in NAFLD, involving an interplay with immune cells (Figure 6). Our key findings are: 1. NAFLD endothelial cells were activated and enriched for chemokine hallmarks and human leukocyte antigens; 2. NAFLD endothelial cells may preferentially recruit immune cells via the CXCL12-CXCR4 axis; 3. NAFLD patients had more pronounced endothelial injury, which may in part be due to heightened interactions with effector immune cells.

**Figure 6:**
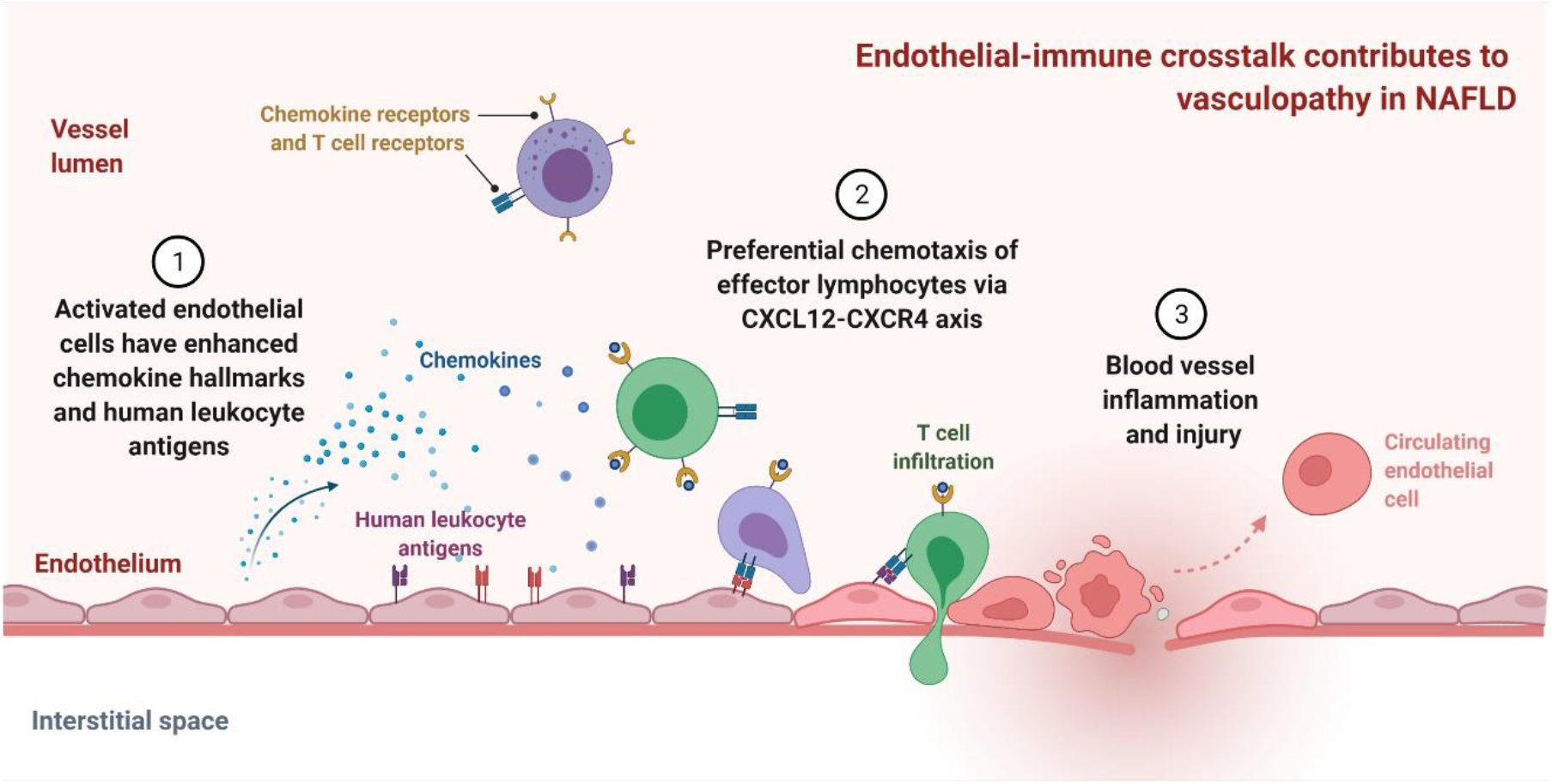
Mechanisms of vascular injury due to NAFLD endothelial chemokine milieu and preferential interaction with effector immune cells via CXCL12-CXCR4 axis.

Several studies have reported elevated chemokines in the plasma/sera of NAFLD patients, as well as the role of these chemokines in regulating liver inflammation (Pan et al., 2020, Marra and Tacke, 2014). Sources of cells capable of contributing to soluble CXCL12 in circulation include cholangiocytes and liver sinusoidal endothelial cells (Mendt and Cardier, 2012, Bigorgne et al., 2008). Here, transcriptomic analysis demonstrated enriched chemokine signatures in NAFLD patient endothelial cells, suggesting that activated endothelial cells could be a potential source of elevated chemokines in NAFLD patients. This was supported by increased *in vivo* expressions of *CXCL12* associated with arterial endothelial cells and liver vasculatures in our mouse NAFLD models. Although CXCL12 has previously been studied for their beneficial impact on atherosclerotic plaque stability and neurovascular repair after stroke, these mainly relate to cell-specific atheroprotective effects and CXCL12’s role in recruitment of bone marrow-derived cells to sites of injury (Zernecke and Weber, 2014, Wang et al., 2012). Instead, plasma CXCL12 levels predict long-term outcomes for stroke and adverse cardiovascular events (Schutt et al., 2012, Ghasemzadeh et al., 2015). Mechanistically, CXCL12 is known to disrupt cholesterol efflux from macrophages by activating GSK-3β/β-catenin/TCF21 signaling pathway, thus exacerbating atherosclerosis (Gao et al., 2019). Notably, endothelial-derived CXCL12 is a key driver of atherosclerosis and one of the contributors to plasma CXCL12 levels (Döring et al., 2019). Atherosclerosis *Apoe*-/-mice deficient of CXCL12 from arterial endothelial cells, but not from smooth muscle cells, bone marrow–derived hematopoietic and nonhematopoietic resident cells, led to marked reduction of lesion size (Döring et al., 2019). Given that chemokine secretions by endothelial cells could influence the recruitment of distinct immune populations (Øynebraåten et al., 2004), we further postulated that NAFLD endothelial cell-derived chemokines could act in an autocrine manner to recruit specific types of immune cells, possibly leading to immune-mediated endothelial damage.

While immune landscape of NAFLD is not yet a well-studied area, there are indications of immune cell population changes and their contribution to NAFLD progression. Alterations of cytotoxic CD8+ T cells are associated with steatohepatitis phenotypes in patients, and CD4+ T helper cells (Th1 and Th17) are positively correlated with severity of NAFLD (Haas et al., 2019). In our previous work using humanized mice reconstituted with human immune components, we found that CD4+ T cells promoted liver fibrosis (Her et al., 2020a). Here, our single-cell analysis pointed to an overarching T cell intensification in NAFLD patients. Correspondingly, our coculture assays using patient-derived endothelial and immune cells revealed preferential chemotaxis of immune subsets including CD4+ T cells, NK cells and monocytes, as well as increased endothelial permeability after immune cell transmigration. Endothelial cells are known to be one of the primary targets of immunologic attack. Effector CD4+ T cells were shown to cause damages to endothelial cells in acute coronary syndrome and systemic sclerosis (Maehara et al., 2020, Nakajima et al., 2002). Endothelial injury also occurs as a result of immune response to infections, graft-versus-host disease and after therapy with immune-activating cytokines (Claser et al., 2019, Chioh et al., 2021, Baik et al., 2021). Autoimmune mechanisms in atherosclerosis involves the production of Th1 cytokines, supporting the inflammatory response that drives endothelial dysfunction (Shoenfeld et al., 2001). The balance between pathogenic T helper cells and beneficial T regulatory cells is tightly controlled by their cytokine environment, and will determine the overall immune response (Leung et al., 2010). Further studies will be needed to elucidate how T cell subsets regulate inflammation in NAFLD setting.

We believe that activated endothelial cells in NAFLD may be providing a chemokine milieu that preferentially interplays with effector immune cells through chemokine ligand-receptor interactions, potentially leading to inflammatory pathology and cytotoxicity-induced vascular injury. Interestingly, inhibition of CXCL12-CXCR4 axis modulated endothelial chemotaxis of immune cells in our coculture assays, and restored endothelial barrier integrity. Current therapeutic exploration on the intervention of chemokines and their receptors in NAFLD are mainly targeting hepatocellular carcinoma (Chen et al., 2018). The recent outcomes of the Canakinumab Anti-inflammatory Thrombosis Outcome Study (CANTOS) prove that immunomodulatory treatment can be used to treat atherosclerosis by reducing inflammatory burden. However, such treatment that use monoclonal antibody targeting interleukin-1β is impeded by its high price, failure to improve overall mortality and an increased risk of infection (Ridker et al., 2017, Sehested et al., 2019). Furthermore, limited success of the Cardiovascular Inflammation Reduction Trial (CIRT) that tested low-dose methotrexate (Ridker et al., 2019), has shown that broad anti-inflammatory treatments are ineffective in reducing cardiovascular events. Therefore, new affordable therapeutics tailored to specific autoimmune mechanisms and with limited immunosuppressive effect are still required. Currently, CXCR4 inhibitor, AMD3100, is used clinically for the treatment of HIV infection and as a hematopoietic stem cell (HSC) mobilizer in non-Hodgkins lymphoma (NHL) patients receiving radiation therapy (Keating, 2011, De Clercq, 2005). More recently, experimental therapeutics using AMD3100 have been explored in Alzheimer’s disease (AD) (Gavriel et al., 2020), Amyotrophic Lateral Syndrome (ALS) (Rabinovich-Nikitin et al., 2016), and Hepatopulmonary Syndrome (HPS) (Shen et al., 2018). In these conditions, AMD3100’s interference with the CXCR4-CXCL12 signaling axis proves useful in reducing inflammatory burden, resulting in attenuated disease progression *in vivo*. Collectively, these findings present modulation of CXCR4-CXCL12 axis as a viable avenue of anti-inflammatory therapeutics.

In evaluating vascular injury in NAFLD, this is the first report of a significantly higher level of circulating endothelial cells (CECs) in NAFLD patients than healthy controls. CECs are not to be confused with endothelial progenitor cells (EPCs) which originate from bone marrow and are recruited to sites of vascular injury to undergo repair (Hebbel, 2017). An altered status of circulating EPCs hence reflects one’s endothelial repair capacity. NAFLD patients were found to have reduced levels of circulating EPCs and impaired adhesive and migratory functions than their healthy counterparts. This suggests that the attenuation of EPC-mediated endothelial repair may contribute to atherosclerotic disease progression and enhanced cardiovascular risk and events in NAFLD patients (Chiang et al., 2012, Francque et al., 2016). CECs, on the other hand, are directly indicative *in situ* vascular injury as they are cells dislodged from damaged endothelia into the bloodstream (Hebbel, 2017). A limitation of this study is that we have not distinguished NAFLD patients with and without confounding cardiovascular risks such as hypertension and diabetes which may also contribute to more CECs (Chioh et al., 2021). CECs are a representative marker of endothelial dysfunction and are useful to serve as a form of vascular health surveillance in conditions that predispose patients to vascular complications.

We present the molecular underpinning of endothelial dysfunction in NAFLD, potentially involving vascular inflammation mediated by effector immune cells. Vascular instability may be further exacerbated by shared cardiometabolic risk profiles common to most NAFLD patients. While there is much complexity in the pathogenesis of NAFLD, intervening specific pathways of endothelial-immune crosstalk may be effective at reducing burdens of vasculopathy in NAFLD patients. Finally, it is imperative to monitor vascular health risk of thromboembolism as part of the management of NAFLD patients so as to ensure early intervention with preventive therapies.

## Materials and Methods

### Study approvals, patient enrolment and sample collection

This study was approved by the Local Ethics Committee of National Healthcare Group Domain Specific Review Board (DSRB Ref: 2016/00580) and Nanyang Technological University Singapore Institutional Review Board (IRB-2020-05-012 and IRB-2020-09-011). Written informed consent was obtained from each participant after the nature and possible consequences of the studies have been explained. The study protocol complies with the Helsinki Declaration.

Patients with biopsy proven NAFLD were taken from the fatty liver clinic, or from the bariatric surgery clinic. Patients with NAFLD had at least 5% steatosis characterized on liver biopsies. NAFLD patients were selected across the spectrum of disease. Controls were selected from healthy volunteers having a Controlled Attenuation Parameter score of less than 248 on vibration controlled transient elastography, indicative of less than 5% steatosis. In addition, control patients did not have raised liver enzymes or evidence of fat in the liver on any other imaging modality.

For sample collection, 10 ml of fresh blood was collected from each participant and processed in the laboratory within 6h. Upon Ficoll centrifugation of the blood specimen, a buffy coat layer containing peripheral blood mononuclear cells (PBMCs) were isolated. PBMC fractions were used for three purposes - (1) Cultivated in cell culture to derive blood outgrowth endothelial cells (BOECs); (2) Immunophenotyping and isolation of immune subpopulations; (3) Analysis of circulating endothelial cells.

### Derivation and maintenance of blood outgrowth endothelial cells

We employed established protocols (14, 15) to derive our BOECs. Peripheral blood samples were processed by Ficoll centrifugation to derive PBMC fraction. Then, PBMCs were seeded onto collagen I-coated wells at a cell density of 1.5 - 2 × 10^6^ cells/cm^2^ in endothelial growth media 2 (EGM-2) medium (Lonza) supplemented with 16% defined fetal bovine serum (FBS; Hyclone).

Outgrowth colonies should appear between 7 to 14 days post-seeding. Quality controls on cell purity, endothelial marker expressions and functional characterization were performed on BOECs from passages 2 and 3 (Supplemental Methods). Only BOEC lines that passed quality control checks would be utilized for downstream experimentations.

### BOEC RNA-sequencing and analysis

Total RNA was isolated from the BOECs. PolyA library preparation and 150 base-pair paired end sequencing was performed by Novogene sequencing facility (Singapore) using an Illumina HiSeq sequencer. The average sequencing depth was 50 million reads. Reads were aligned to the human reference genome GRCh37 using STAR version 2.6.1d. Transcript abundances were quantified with the Partek E/M algorithm based on Ensembl Transcripts Release 83, subsequently normalized by CPM (counts per million) method. Differentially expressed genes in the transcriptome data were identified using the gene specific analysis (GSA) method for *p*-values and Benjamini-Hochberg method for FDR. Details on downstream enrichment and network analyses were described in Results and Supplemental Methods.

### Treatment of BOECs

BOECs were grown to confluence in EGM-2 media supplemented with 10% of their autologous plasma or NAFLD-related risk factors (100 ng/mL of lipopolysaccharide and/or free fatty acids including 100 µM of palmitic acid and 100 µM of oleic acid) for 48h. At the end of treatment, cells were lysed for RNA extraction and qPCR (Supplemental Methods). ELISA of chemokines in plasma and BOEC conditioned media are described in Supplemental Methods.

### Animal studies approval and histological analysis

Animals’ care was in accordance with institutional guidelines, approved by the local Institutional Animal Care and Use Committee (IACUC #: 181367, A18031, A18033). This research complied with the Guidelines on the Care and Use of Animals for Scientific Purposes of the National Advisory Committee for Laboratory Animal Research of Singapore (NACLAR) and the US National Institute of Health (NIH).

Wild-type C57BL/6J male mice (8-9 weeks old) were placed on LIDPAD diet (*n* = 3) or control diet (purified diet containing 14% protein, 76% carbohydrates, 10% fat) for 12 weeks. A related manuscript on this improved LIDPAD diet-induced NAFLD model is in submission elsewhere. After 12 weeks, mice on LIDPAD diet demonstrated histological features representative of the late NASH stage in human NAFLD pathophysiology. Descending thoracic aorta were harvested and aortic specimens were prepared for histological staining (more details in Supplemental Methods).

Humanized mice containing human immune system (HIL, age 32 weeks) were obtained from the Institute of Molecular and Cell Biology, Agency for Science, Technology and Research, Humanized Mouse Unit. Murine models of NAFLD were generated from mice with more than 10% human immune reconstitution (Her et al., 2020b). HIL mice were placed on HFHC diet (Surwit high-fat diet; 58 kcal% fat that is mainly saturated, with carbohydrate-enriched drinking water) or chow diet. After 20 weeks diet, HFHC-HIL mice demonstrated key pathologies representative of human NAFL and NASH, along with visibly increased leukocyte infiltration relative to chow diet-fed mice. Serial liver sections (5 μm thickness) harvested from 4 HFHC-HIL mice (1 male, 3 female) and 4 chow diet-fed mice (2 male, 2 female) were used for immunostaining experiments (more details in Supplemental Methods).

Images were acquired at 40× magnification with a ZEISS Celldiscoverer 7 microscope system. Image analysis was carried out with Fiji software, on the delimitated vascular endothelia of murine aortae and liver sections (Schindelin et al., 2012). At least three sections per animal, with five views per section, were analysed (more details in Supplemental Methods).

### Immunoprofiling by single-cell RNA-sequencing

NAFLD PBMCs were pooled from 2 NAFLD patients while healthy PBMCs were pooled from 4 healthy individuals with each individual contributing about 200,000 cells each (More details in Supplemental Methods). Cell suspensions a final concentration of ∼1200 cells/µl were then loaded onto 10X Genomics Chromium Controller chip by facility personnel at Single-cell Omics Centre (SCOC), Genome Institute Singapore (GIS). NAFLD and healthy PBMC pools were prepared as separate scRNA-seq libraries using Chromium Single Cell 3’ v3 Reagent Kit (10X Genomics) by SCOC GIS and the final ready-to-sequence libraries were handed over with quantification and quality assessment reports from Bioanalyzer Agilent 2100 using the High Sensitivity DNA chip (Agilent Genomics). NAFLD and healthy PBMC libraries were pooled equimolarly and sent for sequencing by NovogeneAIT Genomics (Singapore). Raw sequencing data was also processed by NovogeneAIT Genomics (Singapore) using CellRanger (10x Genomics) with reads mapped to the human genome assembly (GRCh38).

We performed secondary analysis on filtered matrix files using Seurat (v 3.2.0) (Stuart et al., 2019). Data was filtered for dead/poor quality cells based on low number of genes detected (<500) or potential doublets (>6000). Cells with high percentage of mitochondrial genes were also removed at a threshold of less than 35%. These filtered datasets were then scaled and normalized using *SCTransform* individually before integrated based on 3000 integration features. Clusters were identified in the integrated dataset using the *FindCluster* function at resolution 0.3, after PCA analysis, *RunUMAP* and *FindNeighbours* at 1:30 dimensions. Cell type annotation was achieved using DICE annotation (Schmiedel et al., 2018). Differential expression analysis between NAFLD and healthy datasets (RNA) was performed using *FindMarkers* with MAST (Finak et al., 2015) (R package) for each individual cell type. Gene enrichment analysis was carried out for differential expression genes between PCV and normal in each clusters using clusterProfiler (R package, v 3.17.0.) (Yu et al., 2012).

### Endothelial-leukocyte chemotaxis assay

CD8+/-immune cell fractions were isolated by magnetic-activated cell sorting (130-045-201, CD8 MicroBeads, Miltenyi Biotec). Immune cells were allowed to rest for 24h, then labeled with a Deep Red CellTracker (Invitrogen) prior to chemotaxis assay. The assays were performed on 96-well Transwell plates with a membrane pore size of 5 µm, as per the manufacturer’s protocols (CBA-105, Cell BioLabs). Briefly, BOEC monolayers were cultivated in EGM-2 with 2% FBS for 48h in the lower chamber before the CD8+/-cell fractions from top chamber were allowed to migrate for 4h. Then, migrated cells were dislodged and collected from the underside of transwell membranes. The numbers of migrated cells were determined using ZEN 2 software (Zeiss).

### Transendothelial migration assay

BOECs derived from three NAFLD subjects were pooled and seeded to confluence on the membranes with pores of size 5 µm (Corning) of the upper chambers of transwells for 24h. Then, the cells were treated with or without LPS and FFAs for another 48h. In parallel, PBMCs were isolated from blood samples of NAFLD subjects and allowed to rest in RPMI 1640 supplemented with 16% FBS for 24h. On the day of assay, the PBMCs were resuspended in assay buffer (EGM-2 + RPMI 1640 1:1 with 2% FBS and 0.5% BSA) and added to the BOEC monolayers in the upper chambers and the PBMCs were allowed to transmigrate for 16h. BOEC barrier integrity was characterized by transendothelial electrical resistance (TEER).

### Inhibition of CXCL12-CXCR4 axis in chemotaxis by chemokine receptor antagonist

Plerixafor (AMD3100) is a CXCR4 antagonist, reconstituted as per the manufacturer’s instructions (HY-10046, MedChemExpress). Prior to coculture assays (chemotaxis or transendothelial migration assay), the immune cell fractions were pre-incubated with 10 µM of AMD3100 for 30 minutes (Biasci et al., 2020). Assays were then performed as per described above with and without AMD3100.

### Profiling of circulating endothelial cells in NAFLD patients and healthy subjects

CECs were detected from PBMCs through the combined immunophenotypic profile of CD45-/CD31+/CD133-/DNA+ (Supplemental Methods). Flow Cytometry was performed using BD LSRFortessa X-20 (BD Biosciences) and FACSDiva software (BD Biosciences) and data analyzed using FlowJo v10.7.1 software (Becton Dickinson). Each analysis included at least 30,000 cells per individual.

### Statistical analysis

Statistical significance of differences between the cohorts were analyzed using GraphPad Prism version 9. Data normality was determined by Shapiro-Wilk test. Datasets with normal distributions were analyzed with unpaired Student’s two-tailed *t*-tests to compare two conditions; one-or two-way analysis of variance (ANOVA) followed by post-hoc Tukey for datasets with more than two conditions. Non-parametric Mann–Whitney *t*-test or Kruskal-Wallis test were used for non-normally distributed data. Application of these statistical methods to specific experiments is noted in the figure legends. A *p*-value of less than 0.05 were considered significant. Results are depicted as either mean ± standard deviation (SD) or box plots indicating median (middle line), 25th, 75th percentile (box) and the lowest (respectively highest) data points (whiskers).

### Availability of Data and Materials

Further information and requests for resources and reagents should be directed to and will be fulfilled by the corresponding author. Some materials used in this study are commercially procured. There are restrictions to the availability of blood outgrowth endothelial cell lines derived from human patients and normal donors due to ethics considerations for use of these materials within the current scope of study. Requests can be made to the corresponding author as we will explore use of materials subject to new ethics approval and research collaboration agreement (including material transfer).

The authors declare that all data supporting the findings of this study are available within the paper and supplemental materials. Specifically, RNA-sequencing and single-cell sequencing dataset that support the findings of this study will be made publicly available by the publication of this study.

## Acknowledgments

We thank all patients and healthy donors who have participated in this study. Special thanks to EMULSION consortium for their support, Ms Nur Halisah Jumat and the team at National University Hospital for coordinating clinical sample collection, Dr Shyam Prabhakar and Dr Samydurai Sudhagar for advice on 10x Genomics single cell library preparations, Ms Nguyen Le Uyen Nhi and Mr Ryan Gwee Meng Ferng for experimental assistance, Dr Anthony Siau for critical discussion on literature review, and Dr Balakrishnan Kannan for training on image acquisition using the ZEISS Celldiscoverer7 microscope system, Dr Zhisheng Her and Dr Joel Heng Loong Tan for help rendered in humanized mouse liver sections.

This work is funded by an IAF-PP grant (H18/01/a0/017) from the Agency for Science, Technology and Research (Singapore) on Ensemble of Multi-disciplinary Systems and Integrated Omics for NAFLD (EMULSION) diagnostic and therapeutic discovery. The team from Nanyang Technological University Singapore was also funded by the Nanyang Assistant Professorship and Human Frontier Science Program (RGY0069/2019).

## Author Contributions

Conceptualization: CC

Data curation: CYN, KLL, MDM, KXW, FWJC, KT, GSST

Formal analysis: CYN, KLL, MDM, KXW, FWJC, CC

Investigation: CYN, KLL, KXW, FWJC, KT, GSST

Methodology: CYN, KLL, KXW, FWJC, KT, ZSL

Resources: MDM, AS, LWM, QFC, NST

Validation: CYN, KLL

Visualization: CYN, KLL, KXW, FWJC, KT

Funding acquisition: CC, DYY, NHH

Project administration: CC, DYY, NHH

Supervision: CC, DYY, NHH, QFC, NST

Writing – original draft: CC, CYN, KLL

Writing – review & editing: All authors

All authors approved the submitted manuscript version.

## Competing Interest Statement

The authors declare no competing interests.

## Notes

### Competing Interest Statement

The authors have declared no competing interest.

