## Supplemental for "Endothelial-immune crosstalk contributes to vasculopathy in non-alcoholic fatty liver disease"

**This PDF file includes:**

Supplemental Figures S1 to S3 with appropriate legends
Supplemental Tables S1 to S5
Supplemental Materials & Methods
Supplemental References

### Supplementary Information

#### Supplemental Figures and Legends

**Fig. S1**

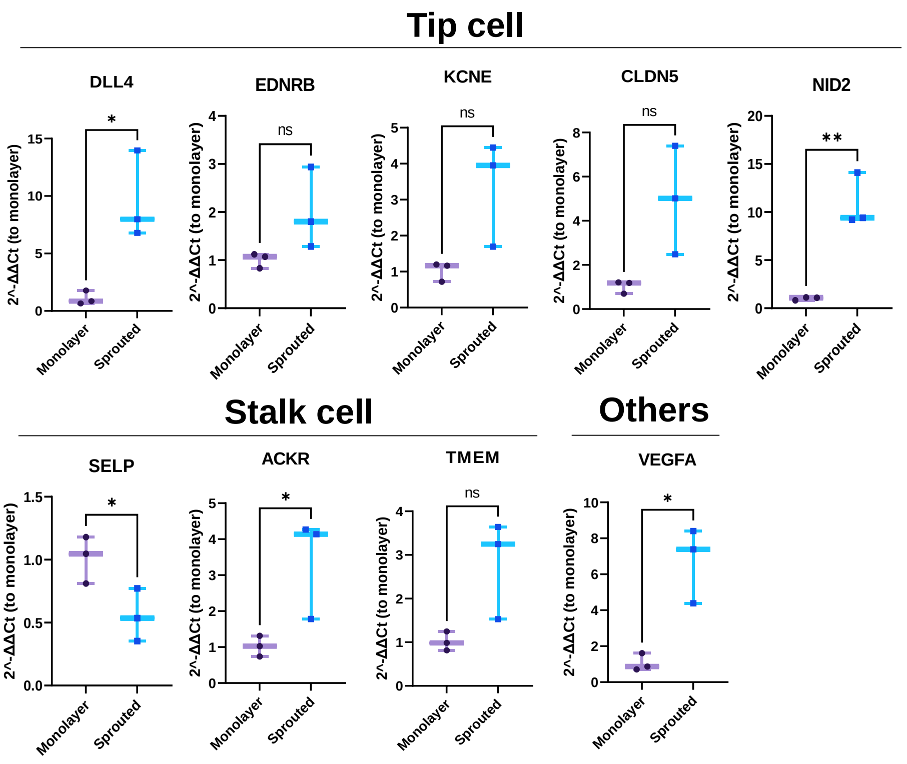


**Fig. S1. Gene expressions of tip-stalk markers in sprouting BOECs.** Expressions were determined by qPCR and normalized to monolayers (non-sprouting). Data presented as mean ± SD. $n = 3$; ns, not significant; *$p < 0.05$; **$p < 0.01$ (*t*-test).

**Fig. S2**

**a**

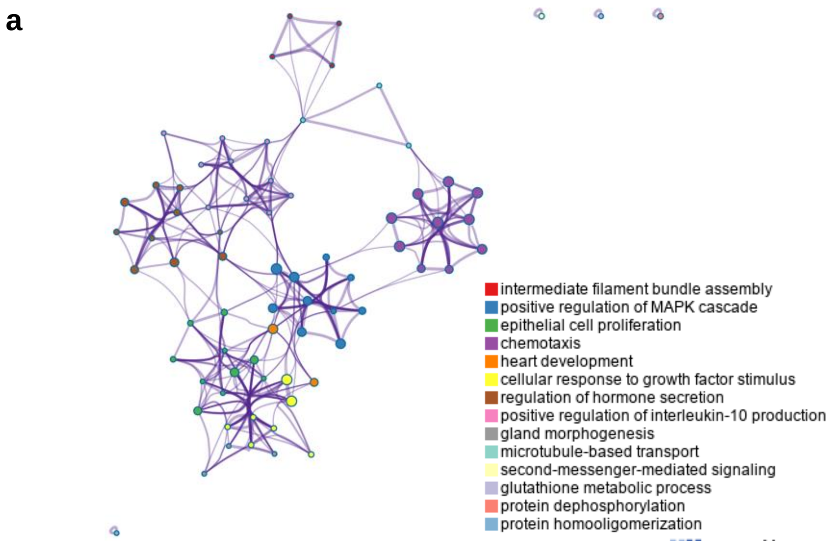


**b**

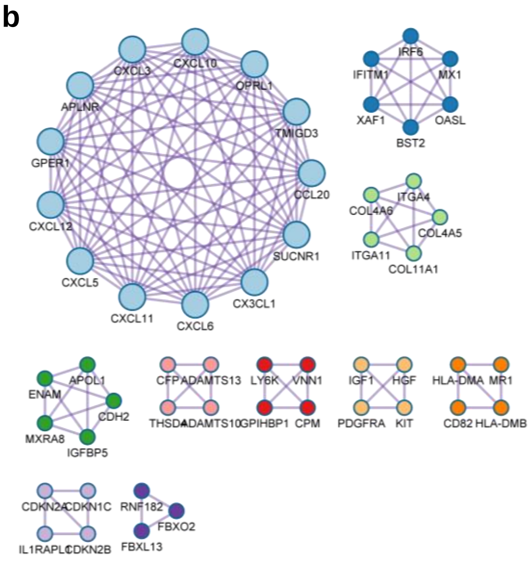


| Color | MCODE | GO | Description | $Log_{10}$(*p*-value) |
|---|---|---|---|---|
| | MCODE_1 | GO:0070098 | chemokine-mediated signaling pathway | -17.1 |
| | MCODE_1 | GO:1990868 | response to chemokine | -16.7 |
| | MCODE_1 | GO:1990869 | cellular response to chemokine | -16.7 |
| | MCODE_2 | GO:0060337 | type I interferon signaling pathway | -14.9 |
| | MCODE_2 | GO:0071357 | cellular response to type I interferon | -14.9 |
| | MCODE_2 | GO:0034340 | response to type I interferon | -14.7 |
| | MCODE_3 | GO:0030198 | extracellular matrix organization | -9.3 |
| | MCODE_3 | GO:0043062 | extracellular structure organization | -9.3 |
| | MCODE_3 | GO:0045229 | external encapsulating structure organization | -9.3 |
| | MCODE_4 | GO:0043687 | post-translational protein modification | -9.5 |
| | MCODE_5 | GO:0030198 | extracellular matrix organization | -5.0 |
| | MCODE_5 | GO:0043062 | extracellular structure organization | -5.0 |
| | MCODE_5 | GO:0045229 | external encapsulating structure organization | -5.0 |
| | MCODE_7 | GO:0014068 | positive regulation of phosphatidylinositol 3-kinase signaling | -10.0 |
| | MCODE_7 | GO:0014066 | regulation of phosphatidylinositol 3-kinase signaling | -9.4 |
| | MCODE_7 | GO:0014065 | phosphatidylinositol 3-kinase signaling | -9.1 |
| | MCODE_8 | GO:0019884 | antigen processing and presentation of exogenous antigen | -6.0 |
| | MCODE_8 | GO:0048002 | antigen processing and presentation of peptide antigen | -5.9 |
| | MCODE_8 | GO:0019882 | antigen processing and presentation | -5.6 |
| | MCODE_9 | GO:0045736 | negative regulation of cyclin-dependent protein serine/threonine kinase activity | -8.3 |
| | MCODE_9 | GO:1904030 | negative regulation of cyclin-dependent protein kinase activity | -8.2 |
| | MCODE_9 | GO:0000079 | regulation of cyclin-dependent protein serine/threonine kinase activity | -6.7 |

**Fig. S2. Transcriptomes of healthy and NAFLD BOECs. (a)** Metascape enrichment network visualization of interactomes of the down-regulated terms in NAFLD BOECs. No identifiable MCODE complexes were found. **(b)** Ten MCODE complexes identified in the up-regulated networks in NAFLD BOECs, ranked and colored by their identities. The identities of the complexes were listed in the table.

Fig. S3

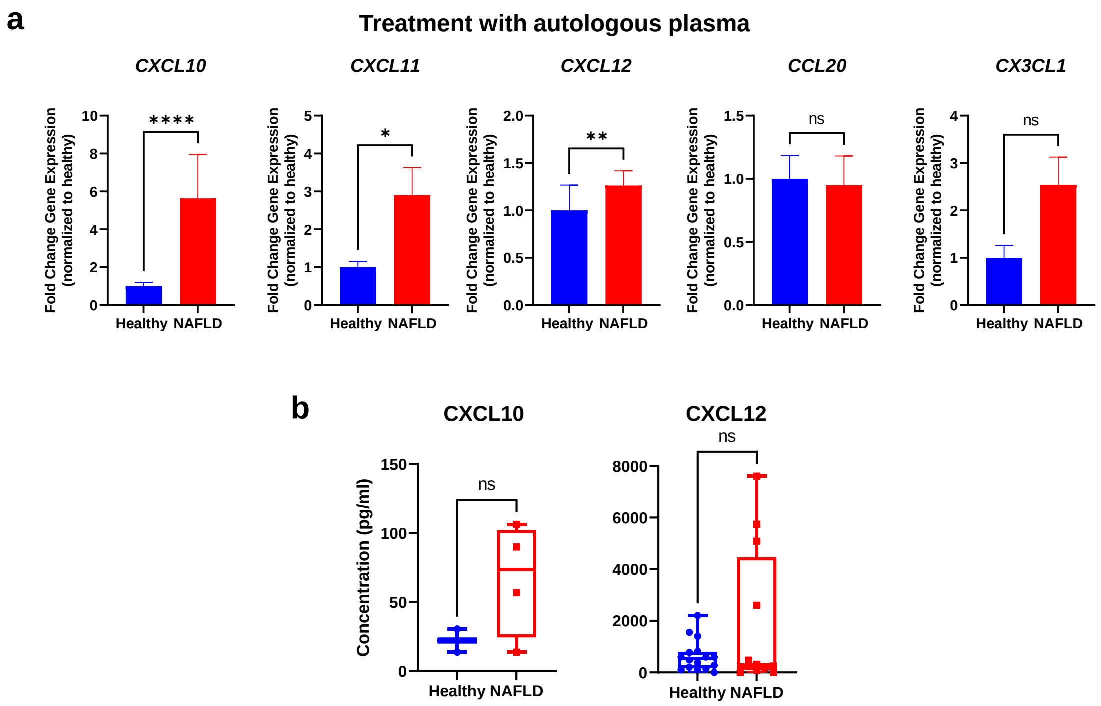


**Fig. S3. (a)** Chemokine expressions in the BOECs treated with autologous plasma. Fold changes in gene expressions of *CXCL10*, *CXCL11*, *CXCL12*, *CCL20* and *CX3CL1* in NAFLD ($n$ = 11) BOECs were normalized to healthy ($n$ = 12) BOECs. ns, not significant; *$p$<0.05; **$p$<0.01; ****$p$<0.0001 (Mann-Whitney test). Results are indicated by mean ± SD. **(b)** Plasma concentrations of CXCL10 and CXCL12 in healthy and NAFLD subjects. ns, not significant (Mann-Whitney test).

**Supplemental Tables**

**Table S1.** Baseline characteristics of healthy controls and NAFLD patients whose BOEC lines were subjected to RNA-sequencing.

| Diagnosis | Gender | Age | Ethnicity | Comorbidities | | |
|---|---|---|---|---|---|---|
| | | | | Diabetes | Hypertension | Dyslipidemia |
| NAFLD (steatosis) | Female | 31 | Asian | No | No | No |
| | Female | 51 | Asian | No | Yes | No |
| | Male | 31 | Asian | Yes | Yes | Yes |
| Healthy | Female | 51 | Asian | No | No | No |
| | Female | 34 | Asian | No | No | No |
| | Male | 40 | Asian | No | No | No |

**Table S2.** Top genes prioritized using 12 topological analysis methods on the significantly upregulated genes in NAFLD BOECs.

Methods

Genes

| MCC | DMNC | MNC | Deg | EPC | BN | EC | CLO | Rad | BC | STR | CC |
|---|---|---|---|---|---|---|---|---|---|---|---|
| CXCL10 | CX3CL1 | CX3CL1 | CXCL12 | CX3CL1 | CXCL10 | CXCL10 | CXCL10 | CXCL10 | CXCL10 | CXCL10 | CX3CL1 |
| CXCL12 | OPRL1 | OPRL1 | CXCL10 | SUCNR1 | OASL | CX3CL1 | CXCL12 | MX1 | IL33 | CXCL12 | OPRL1 |
| GPER1 | SUCNR1 | SUCNR1 | IGF1 | GPER1 | IL33 | OPRL1 | GPER1 | OASL | GATA3 | IGF1 | IL1RL1 |
| CCL20 | CXCL5 | CXCL5 | GPER1 | CXCL12 | CXCL12 | SUCNR1 | MX1 | CXCL12 | BST2 | BST2 | SUCNR1 |
| CX3CL1 | CXCL3 | CXCL3 | CCL20 | CXCL6 | GATA3 | IL33 | CCL20 | GPER1 | CXCL12 | IL33 | COL4A6 |
| OPRL1 | GPER1 | GPER1 | CX3CL1 | APLNR | BST2 | CXCL5 | OASL | IL33 | HIST1H2BJ | ATP8A1 | FOSB |
| SUCNR1 | CXCL12 | CXCL12 | OPRL1 | CXCL11 | HIST1H2BJ | CXCL3 | CX3CL1 | IFI44L | ATP8A1 | GATA3 | CXCL5 |
| CXCL5 | ADORA3 | ADORA3 | SUCNR1 | CXCL5 | ATP8A1 | MX1 | OPRL1 | CCL20 | SNAP25 | MX1 | CXCL3 |
| CXCL3 | CXCL10 | CXCL10 | CXCL5 | CXCL3 | COL11A1 | GPER1 | SUCNR1 | CX3CL1 | MX1 | OASL | MXRA8 |
| ADORA3 | CXCL6 | CXCL6 | CXCL3 | OPRL1 | TLL1 | IFI44L | CXCL5 | OPRL1 | OASL | SNAP25 | CEACAM1 |
| CXCL6 | APLNR | APLNR | ADORA3 | CXCL10 | SNAP25 | CXCL12 | CXCL3 | SUCNR1 | IGF1 | HIST1H2BJ | CPM |
| APLNR | CCL20 | CCL20 | CXCL6 | ADORA3 | HGF | ADORA3 | ADORA3 | CXCL5 | CDK4 | HGF | ENAM |
| CXCL11 | CXCL11 | CXCL11 | APLNR | CCL20 | IGF1 | OASL | CXCL6 | CXCL3 | HGF | KITLG | TNFRSF10B |
| IGFBP5 | MXRA8 | MX1 | CXCL11 | MX1 | HERC5 | CXCL6 | APLNR | ADORA3 | HERC5 | FGF13 | ADAMTS13 |
| IGFBP1 | ENAM | OASL | SNAP25 | OASL | CFP | APLNR | CXCL11 | CXCL6 | KITLG | GPER1 | ADORA3 |
| IGFBP4 | APOL1 | XAF1 | MX1 | XAF1 | CDK4 | CCL20 | IFI44L | APLNR | VNN1 | IGFBP5 | TNFSF10 |
| MXRA8 | CDH2 | IGF1 | OASL | IFI44L | EFNB3 | CXCL11 | IGF1 | CXCL11 | FGF13 | IGFBP1 | GPIHBP1 |
| ENAM | BST2 | IFI44L | XAF1 | IGF1 | NOTCH3 | IL1A | HGF | BST2 | GPER1 | IGFBP4 | LIN7A |
| APOL1 | IRF6 | IGFBP5 | IFI44L | KITLG | GRM1 | IL1RL1 | KITLG | HGF | ESR2 | ESR2 | ASAH1 |
| CDH2 | IGFBP5 | IGFBP1 | HERC5 | HGF | KCNN4 | BST2 | FGF13 | KITLG | IFI44L | HERC5 | KCNN3 |

Genes listed are hub genes identified using the 12 topological analysis methods in Cytoscape plugin cytoHubba. Those marked in red indicate chemokines.

Abbreviations are: MCC, Maximum Neighborhood Component; DMNC, Density of Maximum Neighborhood Component; MNC, Maximal Clique Centrality; Deg, Degree; EPC, Edge Percolated Component; BN, BottleNeck; EC, EcCentricity; CLO, Closeness; Rad, Radiality; BC, Betweenness; STR, Stress; CC, ClusteringCoefficient.

**Table S3.** Baseline characteristics of healthy controls (n = 12) and NAFLD patients (n = 11) whose BOEC lines were used for chemokine validation experiments.

| Parameter | Healthy (*n* = 12) | NAFLD (*n* = 11) | *p*-value |
|---|---|---|---|
| **Demographics** | | | |
| Age (year) | 53.5 (40.5 – 61.8) | 35 (28.0 – 39.0) | 0.0010[a, *] |
| Gender (female) | 8 (66.7) | 8 (72.7) | >0.9999[b] |
| Race<br>1) Asian<br>2) Non-Asian | <br>12 (100.0)<br>0 (0) | <br>11 (100.0)<br>0 (0) | <br>>0.9999[c] |
| Weight (kg) | 62.9 (54.4 – 73.2) | 108.2 (104.6 – 117.2) | <0.0001[b, *] |
| BMI ($kg/m^2$) | 23.7 (19.5 – 25.0) | 42.9 (37.8 – 47.5) | <0.0001[a, *] |
| Smoking status<br>1) Never<br>2) Current<br>3) Ever<br>4) Not recorded | <br>11 (91.7)<br>0 (0.0)<br>1 (8.3)<br>0 (0.0) | <br>5 (45.5)<br>4 (36.4)<br>1 (9.1)<br>1 (9.1) | <br>0.0592[c] |
| Alcohol consumption<br>1) Never<br>2) Current<br>3) Ever<br>4) Not recorded | <br>10 (83.3)<br>2 (16.7)<br>0 (0.0)<br>0 (0.0) | <br>8 (72.7)<br>2 (18.2)<br>0 (0.0)<br>1 (9.1) | <br>0.3158[c] |
| **Clinical characteristics** | | | |
| Diabetes mellitus | 0 (0) | 2 (18.2) | 0.2174[d] |
| Hypertension | 0 (0) | 2 (18.2) | 0.2174[d] |
| Dyslipidemia | 0 (0) | 2 (18.2) | 0.2174[d] |
| **Blood lipoproteins** | | | |
| Total cholesterol (mmol/L) | 5.0 (4.1 – 5.5) | 4.9 (4.4 – 6.0) | 0.4205[a] |
| Triglycerides (mmol/L) | 0.8 (0.6 – 1.3) | 0.9 (0.7 – 1.4) | 0.6165[a] |
| **Liver function test and histopathology** | | | |
| ALT (IU/L) | 14.0 (11.5 – 18.5) | 26.0 (19.0 – 42.0) | 0.0045[a, *] |
| AST (IU/L) | 22.0 (19.3 – 24.8) | 22.0 (19.0 – 24.0) | 0.7820[a] |
| Biopsy histopathology[1]<br>1) Steatosis<br>2) Inflammation<br>3) Ballooning<br>4) Fibrosis | <br>N/A | <br>11 (100.0%)<br>2 (18.2%)<br>0 (0.0%)<br>2 (18.2%) | <br>N/A |

Data were presented as median (interquartile range) for continuous variables and *n* (%) for categorical variables.

Abbreviations are: ALT, alanine transaminase; AST, aspartate aminotransferase; BMI, body mass index.

[a]unpaired *t*-test; [b]Mann-Whitney test; [c]Chi-square test for trend; [d]Fisher's exact test; *statistically significant difference $p<0.05$.

[1]Liver histopathology assessment was performed based on the NASH CRN Histologic Scoring System by HistoIndex's AI-based diagnostic technology.

**Table S4.** Baseline characteristics of healthy controls and NAFLD patients whose PBMCs were subjected to immunoprofiling by single-cell RNA-sequencing.

| Diagnosis | Gender | Age | Ethnicity | Comorbidities | | |
|---|---|---|---|---|---|---|
| | | | | Diabetes | Hypertension | Dyslipidemia |
| NAFLD | Female | 58 | Asian | No | No | No |
| NAFLD | Male | 25 | Asian | No | Yes | No |
| Healthy | Female | 46 | Asian | No | No | No |
| Healthy | Male | 47 | Asian | No | No | No |
| Healthy | Female | 60 | Asian | No | No | No |
| Healthy | Male | 43 | Asian | No | No | No |

**Table S5.** Resource table of antibodies, biochemical kits, and primers.

***Antibodies***

| Antibody | Clone | Fluorochrome | Concentration | Manufacturer (catalogue) | Application |
|---|---|---|---|---|---|
| CD31 | WM59 | APC | 100 ng/million cells | BioLegend (303116) | Flow cytometry for BOEC and immune characterizations |
| CD144 | 55-7H1 | PE | 100 ng/million cells | BD Bioscience (421561714) | |
| CD45 | 2D1 | APC | 100 ng/million cells | Invitrogen (17-9459-42) | |
| CD133 | 7 | APC | 100 ng/million cells | BioLegend (372806) | |
| CD68 | Clone: Y1/82A | FITC | 100ng/million cells | Biolegend (333806) | |
| CD8 | SK1 | PE | 5 µL/million cells | BioLegend (344706) | |
| CD4 | RPA-T4 | APC | 100 ng/million cells | BioLegend (300537) | |
| CD45RA | HI100 | APC | 100 ng/million cells | ThermoFisher (17-0458-42) | |
| CCR7 | 3D12 | FITC | 100 ng/million cells | ThermoFisher (11-1979-42) | |
| CDH5 | Polyclonal | Non-conjugated | 4 µg/mL | Santa Cruz (sc-6458) | Histology |
| SDF1 | Polyclonal | Non-conjugated | 5 µg/mL | Bioss (bs-4938R) | |
| Goat IgG (H+L) | Polyclonal | AF568 | 4 µg/mL | Invitrogen (A11057) | |
| Rabbit IgG (H&L) | Polyclonal | AF488 | 10 µg/mL | Abcam (ab150073) | |
| Hoechst 33342 | N/A | N/A | 5 µL/million cells | Invitrogen (R37165) | Flow cytometry for CEC |
| CD31 | WM59 | PE/cyanine 7 | 4 µL/million cells | BioLegend (303118) | |
| CD133 | W6B3C1 | APC | 4 µL/million cells | BioLegend (397906) | |
| CD45 | 2D1 | PE | 3 µL/million cells | BioLegend (368510) | |

***ELISA kits***

| Chemokine | ELISA Kit (Manufacturer, catalogue) |
|---|---|
| CXCL10 | Human IP-10 SimpleStep ELISA Kit (Abcam, ab173194) |
| CXCL11 | I-TAC Human SimpleStep ELISA Kit (Abcam, ab187392) |
| CXCL12 | SDF1-alpha Human ELISA Kit (Abcam, ab100637) |
| CCL20 | Human MIP-3 alpha ELISA Kit (Abcam, ab269562) |
| CX3CL1 | Fractalkine Human SimpleStep ELISA Kit (Abcam, ab192145) |

***Primer sequences for qPCR***

| Gene | Forward primer (5′–3′) | Reverse primer (5′–3′) |
|---|---|---|
| *CXCL10* | 5’-TGCCATTCTGATTTGCTGCC-3’ | 5’-TGCAGGTACAGCGTACAGTT-3’ |
| *CXCL11* | 5’-GGCTTCCCCATGTTCAAAG-3’ | 5’-CAGATGCCCTTTTCCAGGAC-3’ |
| *CXCL12* | 5’-TCAGCCTGAGCTACAGATGC-3’ | 5’-CTTTAGCTTCGGGTCAATGC-3’ |
| *CCL20* | 5’-ATGTGCTGTACCAAGAGTTTG-3’ | 5’-TTACATGTTCTTGACTTTTTTACTGAGGAG-3’ |
| *CX3CL1* | 5’-ATGGCTCCGATATCTCTG-3’ | 5’-TGCTGCATCGCGTCCTTG-3’ |

### Supplemental Materials and Methods

#### Derivation and maintenance of blood outgrowth endothelial cells

We employed established protocols (14, 15) to derive our BOECs. Peripheral blood samples (5 – 9 ml) was diluted 1:1 with phosphate-buffered saline and separated to obtain buffy coat by density gradient centrifugation over Ficoll® Paque (GE Healthcare). The buffy coat, which was enriched with PBMCs, was carefully collected, washed with PBS, resuspended in endothelial growth media 2 (EGM-2) medium (Lonza) supplemented with 16% defined fetal bovine serum (FBS; Hyclone). Then, the PBMCs were seeded onto collagen I-coated wells at a cell density of 1.5 - 2 × $10^6$ cells/$cm^2$. Culture medium was changed every two to three days. Outgrowth colonies should appear between 7 to 14 days post-seeding. Quality controls on cell purity, endothelial marker expressions and functional characterization were performed on BOECs from passages 2 and 3. Only BOEC lines that passed quality control checks would be utilized for downstream experimentations.

#### Endothelial marker characterization by flow cytometry

Cell surface markers were quantified using flow cytometry to phenotypically confirm endothelial identity of the derived BOECs. Briefly, BOECs were washed and stained with a cocktail of surface antibodies in staining buffer containing PBS with 2% FBS, for 15 minutes in the dark. CD31 (platelet endothelial cell adhesion molecule 1; clone wm59, BioLegend) and CD144 (vascular endothelial cadherin; clone 55-7H1, BD Pharmingen) were selected as endothelial markers, CD45 (clone 2D1, Invitrogen) as leukocyte exclusion marker, and CD133 (clone 7, BioLegend) as a progenitor marker. Fluorescence data were collected on a BD LSR Fortessa X-20 cell analyzer (Becton Dickinson) and analyzed using FlowJo v10.7.1 software (Becton Dickinson). Details on antibodies are included in Supplemental Table S5.

#### Endothelial marker characterization by immunocytochemistry

Confluent monolayers of BOECs were fixed and stained with anti-CD31 (clone 89C2, 1:200, Cell Signaling) and anti-CD144 (clone C-19, 1:200, Santa Cruz) for identity verification. Staining was detected with secondary antibodies Alexa-Fluor 488 and 568 and visualized using ZEISS Celldiscoverer7 microscope system. All experiments were accompanied by a negative control – fibroblasts. Details on antibodies are included in Supplemental Table S5.

#### Endothelial functional characterization by angiogenesis assays

Fibrin gel bead-based sprouting angiogenesis assay was performed as per described (1). Briefly, BOECs were coated onto collagen-coated Cytodex 3 microcarrier beads (Sigma-Aldrich) at 150 cells/bead for 4h and allowed to adhere overnight. Then, the coated beads were suspended in a 2.0 mg/ml of fibrinogen solution (Sigma-Aldrich) at a concentration of 500 beads/ml. Fibrin gels were formed by adding 500 µL of the fibrinogen/bead suspension to each well of a 24-well plate containing 0.625 U/mL of thrombin (Sigma-Aldrich). Once clotted, the gels were topped with 0.5 ml of EMG-2 supplemented with 10% heat-inactivated FBS (Gibco). Cells were incubated at 37°C/5% $CO_2$ and sprouting was observed after 24h using bright-field microscopy (Lumenera).

#### BOEC RNA-sequencing and analysis

Total RNA was isolated from the BOECs as per manufacturer's instructions (RNeasy Micro Kit, Qiagen). PolyA library preparation and 150 base-pair paired end sequencing was performed by Novogene sequencing facility (Singapore) using an Illumina HiSeq sequencer. The average sequencing depth was 50 million reads. All subsequent analyses were performed on Partek Flow software, version 7.0 (St. Louis, MO, USA). Reads were aligned to the human reference genome GRCh37 using STAR version 2.6.1d. Transcript abundances were quantified with the Partek E/M algorithm based on Ensembl Transcripts Release 83, subsequently normalized by CPM (counts per million) method. Differentially expressed genes in the transcriptome data were identified using the gene specific analysis (GSA) method for *p*-values and Benjamini-Hochberg method for FDR. Genes were called differentially expressed if $p$-value < 0.05, FDR < 0.05, and fold change ≥ 2 or ≤ -2. Enrichment analyses for differentially expressed genes were performed using Metascape (2). Protein-protein interactions were analyzed using the STRING tool, and the yielded networks were further constructed and analyzed in Cytoscape (version 3.8.0) with the network topography based on the confidence score calculated by STRING. Cytoscape plug-in cytoHubba was used to assess the centrality (strongly interconnected subnetworks) of differentially expressed genes in the protein-protein interaction networks with 12 algorithms: Maximum Neighborhood Component (MCC), Density of Maximum Neighborhood Component (DMNC), Maximal Clique Centrality (MNC), Degree (Deg), Edge Percolated Component (EPC), BottleNeck (BN), EcCentricity (EC), Closeness (CLO), Radiality (Rad), Betweenness (BC), Stress (STR) and ClusteringCoefficient (CC). The selected hub genes were verified by MCODE method with default algorithms (degree cut-off of 2, node score cut-off of 0.2, K-Core of 2, and maximum depth of 100).

**Quantitative PCR**
Total RNA was isolated using RNeasy Plus Mini kit (Qiagen) as per the manufacturer's protocol, subsequently used to generate cDNA with LunaScriptTM RT SuperMix Kit (New England Biolabs). Real-time PCR was performed using SYBR green gene expression assays (New England Biolabs) on a QuantStudio 6 instrument (Applied Biosystems). Gene expressions were normalized to endogenous GAPDH housekeeping gene.

**ELISA of chemokines**
EGM-2 media conditioned by confluent monolayers of BOECs were collected after 48h. Secretion of IP-10 (CXCL10, Abcam, ab173194), SDF-1 (CXCL12, Abcam, ab100637), MIP-3 alpha (CCL20, Abcam, ab269562) and fractalkine (CX3CL1, Abcam, ab192145) into conditioned EGM-2 were measured by ELISA as per the manufacturer's protocols. Reads were corrected for background with non-conditioned EGM-2.

**Animal studies approval and histological analysis**
Animals' care was in accordance with institutional guidelines, approved by the local Institutional Animal Care and Use Committee (IACUC #: 181367, A18031, A18033). This research complied with the Guidelines on the Care and Use of Animals for Scientific Purposes of the National Advisory Committee for Laboratory Animal Research of Singapore (NACLAR) and the US National Institute of Health (NIH).
Wild-type C57BL/6J male mice (8-9 weeks old) were placed on LIDPAD diet ($n$ = 3) or control diet (purified diet containing 14% protein, 76% carbohydrates, 10% fat) for 12 weeks. The manuscript relating to this novel LIDPAD diet-induced NAFLD model is in submission elsewhere. After 12 weeks, mice on LIDPAD diet demonstrated histological features representative of the late NASH stage in human NAFLD pathophysiology. We did not use any anaesthetic agents during the course of study or endpoint. Mice were euthanized using $CO_2$. Descending thoracic aorta were then harvested and aortic specimens were fixed overnight in 4% paraformaldehyde (PFA) and embedded in paraffin. Every 1st, 10th and 20th serial sections (5 μm thickness) were mounted onto each slide for histoimmunostaining experiments.
Humanized mice containing human immune system (HIL, age 32 weeks) were obtained from the Institute of Molecular and Cell Biology (IMCB), Agency for Science, Technology and Research, Humanized Mouse Unit. Briefly, human immune reconstitution of NOD-*scid* $Il2r\gamma^{null}$ (NSG) mice was performed via intrahepatic injection of human CD34+ fetal liver cells into one to three-day-old NSG pups following sub-lethal irradiation. Murine models of NAFLD were generated from mice with more than 10% human immune reconstitution, determined according to described criteria (3). We did not use any anesthetic agents during the course of study or endpoint. HIL mice were euthanized using $CO_2$ after being placed on HFHC diet (Surwit high-fat diet; 58 kcal% fat that is mainly saturated, with carbohydrate-enriched drinking water) or chow diet for 20 weeks. After 20 weeks diet, HFHC-HIL mice demonstrated key pathologies representative of human NAFL and NASH, along with visibly increased leukocyte infiltration relative to chow diet-fed mice. Serial liver sections (5 μm thickness) harvested from 4 HFHC-HIL mice (1 male, 3 female) and 4 chow diet-fed mice (2 male, 2 female) were used for immunostaining experiments.

Double immunostaining of CDH5 and CXCL12 was performed on the paraffin-embedded murine aortic and liver sections. Heat induced epitope retrieval (HIER) was performed on deparaffinized sections in the R-Universal epitope recovery buffer (Aptum Biologics, #AP0530-500), using the

2100 Antigen Retriever (Aptum Biologics). Sections were permeabilized with 1X PBS with 0.3% Triton X-100 for 10 minutes and blocked with 1X PBS with 1% BSA for 1h to minimize non-specific staining Sections were incubated in a primary antibody mixture of goat polyclonal anti-CDH5 (Santa Cruz Biotechnology, sc-6458; 4 µg/mL) and rabbit polyclonal anti-CXCL12 (Bioss, bs-4938R; 5 µg/mL) for 1h at room temperature, followed by a secondary antibody mixture of AF568-conjugated donkey anti-goat IgG (Invitrogen, A11057; 4 µg/mL) and AF488-conjugated donkey anti-rabbit IgG (abcam, ab150073, 10 µg/mL) for 1h at room temperature in humidified chambers. All sections were counter-stained with DAPI. Slides were mounted using Fluoromount$^{TM}$ Aqueous Mounting Medium (Sigma-Aldrich, F-4680) in preparation for image acquisition. Details on antibodies are included in Supplemental Table S5.

Images were acquired at 40X magnification with a ZEISS Celldiscoverer 7 microscope system. Image analysis was carried out with Fiji software, on the delimitated vascular endothelia of murine aortae and liver sections (4). For images of murine aorta, vascular endothelia were manually delimitated using region of interest (ROI) drawing tools based on CDH5 signals and internal elastin lamellae. Following background subtraction, CXCL12 fluorescent intensity within the delimitated endothelia was quantified as the sum of pixel intensities in the CXCL12 single-channel image, normalized by the number of pixels with above-zero intensities in the DAPI single-channel image. For images of murine liver, sinusoidal endothelial regions were delimitated by applying a histogram-based automated thresholding on the CDH5 single-channel image. Using the EzColocalization plugin on Fiji, Mander's Coefficient (M2) was obtained as a measure for the degree of CXCL12 overlap with CDH5 within delimitated endothelia (5). M2 indicates the total number of above-threshold CXCL12 pixels as a fraction of above-threshold CDH5 pixels. Threshold values (FT) were set to 0.1 for both CXCL12 and CDH5 channels, meaning that only the 10% of pixels with the highest signals in each channel would be analyzed. For both vascular endothelia of murine aortae and liver sections, there were 10-20 independent regions of interest analyzed per mouse for image quantification.

**Immunoprofiling by single-cell RNA-sequencing**
Cryopreserved PBMCs were thawed in a 37°C water bath and resuspended in Iscove's Modified Dulbecco's Medium (Gibco) supplemented with 10% heat inactivated FBS. PBMCs were then pelleted out of solution by centrifuging at 300g for 5 minutes. Cell number and viability percentage were determined using Trypan blue and an automated cell counter (Countess II, Thermo Fisher Scientific). NAFLD PBMCs were pooled from 2 NAFLD patients while healthy PBMCs were pooled from 4 healthy individuals with each individual contributing about 200,000 cells each. Final PBMCs pools were pelleted down at 300g for 5 minutes and resuspended in PBS with 0.04% BSA to give a final concentration of ~1200 cells/µl.
Cell suspensions were then loaded onto 10X Genomics Chromium Controller chip by facility personnel at Single-cell Omics Centre (SCOC), Genome Institute Singapore (GIS). NAFLD and healthy PBMC pools were prepared as separate scRNA-seq libraries using Chromium Single Cell 3' v3 Reagent Kit (10X Genomics) by SCOC GIS and the final ready-to-sequence libraries were handed over with quantification and quality assessment reports from Bioanalyzer Agilent 2100 using the High Sensitivity DNA chip (Agilent Genomics). NAFLD and healthy PBMC libraries were pooled equimolarly and sent for sequencing by NovogeneAIT Genomics (Singapore). Raw sequencing data was also processed by NovogeneAIT Genomics (Singapore) using CellRanger (10x Genomics) with reads mapped to the human genome assembly (GRCh38).

We performed secondary analysis on filtered matrix files using Seurat (v 3.2.0) (6). Data was filtered for dead/poor quality cells based on low number of genes detected (<500) or potential doublets (>6000). Cells with high percentage of mitochondrial genes were also removed at a threshold of less than 35%. These filtered datasets were then scaled and normalized using *SCTransform* individually before integrated based on 3000 integration features. Clusters were identified in the integrated dataset using the *FindCluster* function at resolution 0.3, after PCA analysis, *RunUMAP* and *FindNeighbours* at 1:30 dimensions. Cell type annotation was achieved through comparisons of various iterations from celldex (7) (R package) and known canonical markers of PBMC cell types. We finally settled on DICE annotation (8). Differential expression analysis between NAFLD and healthy datasets (RNA) was performed using *FindMarkers* with MAST (9) (R package) for each individual cell type. Gene enrichment analysis was carried out for differential expression genes between PCV and normal in each clusters using clusterProfiler (R package, v 3.17.0.) (10).

**Immune characterization by flow cytometry**
Immune subsets were separated through MACS (Miltenyi Biotec). Phenotyping of both CD8+/- cell fractions was performed using flow cytometry antibodies directed against CD8-PE (clone SK1, Biolegend 344706), CD4-APC (clone RPA-T4, Biolegend 300537). Details on antibodies are included in Supplemental Table S5. Following 30 minutes of antibody incubation at room temperature, stained cells were washed and suspended in PBS containing 2% heat inactivated FBS. Flow cytometry was performed on BD LSRFortessa$^{TM}$ X-20 Flow Cytometer (BD Biosciences). The gating was set by forward and side scatter, and 10,000 events were acquired. Finally, data were collected and analyzed using FlowJo v10.7.1 software (Becton Dickinson).

**Endothelial-leukocyte adhesion assay**
Adhesion assay was as per described (11). Briefly, THP-1 and Jurkat cells were stained with a Deep Red CellTracker fluorescent dye (Invitrogen). Then, $2.0 \times 10^5$ cells were added on every $cm^2$ of BOEC monolayers for 1h. Subsequently, non-adherent cells were removed, and the number of adherent cells was determined through fluorescence intensity readouts from a Synergy H1 microplate reader (BioTek).

**Profiling of circulating endothelial cells in NAFLD patients and healthy subjects**
100 µl of 1 million PBMCs (whole PBMC fraction from processed blood) per subject were stained in the dark for 10 minutes at room temperature, followed by 20 minutes at 4°C on an analog tube rotator with antibodies (Supplemental Table S5: key resource table). After incubation, cells were washed and resuspended in 200µl of PBS with 1% BSA for flow cytometry analysis. CECs were detected through the combined immunophenotypic profile of CD45-/CD31+/CD133-/DNA+. Details on antibodies are included in Supplemental Table S5. The number of CECs was expressed as cells per million of PBMCs. Flow Cytometry was performed using BD LSRFortessa X-20 (BD Biosciences) and FACSDiva software (BD Biosciences) and data analyzed using FlowJo v10.7.1 software (Becton Dickinson). Each analysis included at least 30,000 cells per individual.

**Supplemental Information References**


1. Nakatsu MN, Davis J, & Hughes CC (2007) Optimized fibrin gel bead assay for the study of angiogenesis. *J Vis Exp* (3):186.
2. Zhou Y, *et al.* (2019) Metascape provides a biologist-oriented resource for the analysis of systems-level datasets. *Nat Commun* 10(1):1523.
3. Her Z, *et al.* (2020) CD4(+) T Cells Mediate the Development of Liver Fibrosis in High Fat Diet-Induced NAFLD in Humanized Mice. *Front Immunol* 11:580968.
4. Schindelin J, *et al.* (2012) Fiji: an open-source platform for biological-image analysis. *Nat Methods* 9(7):676-682.
5. Stauffer W, Sheng H, & Lim HN (2018) EzColocalization: An ImageJ plugin for visualizing and measuring colocalization in cells and organisms. *Sci Rep* 8(1):15764.
6. Stuart T, *et al.* (2019) Comprehensive Integration of Single-Cell Data. *Cell* 177(7):1888-1902 e1821.
7. Aran D, *et al.* (2019) Reference-based analysis of lung single-cell sequencing reveals a transitional profibrotic macrophage. *Nat Immunol* 20(2):163-172.
8. Schmiedel BJ, *et al.* (2018) Impact of Genetic Polymorphisms on Human Immune Cell Gene Expression. *Cell* 175(6):1701-1715.e1716.
9. Finak G, *et al.* (2015) MAST: a flexible statistical framework for assessing transcriptional changes and characterizing heterogeneity in single-cell RNA sequencing data. *Genome Biol* 16:278.
10. Yu G, Wang LG, Han Y, & He QY (2012) clusterProfiler: an R package for comparing biological themes among gene clusters. *OMICS* 16(5):284-287.
11. Wilhelmsen K, Farrar K, & Hellman J (2013) Quantitative in vitro assay to measure neutrophil adhesion to activated primary human microvascular endothelial cells under static conditions. *J Vis Exp* (78):e50677.
